# Programming the Internal Architecture of Synthetic Compartments by Coassembling Filamentous and Liquid DNA Phases

**DOI:** 10.64898/2026.07.20.739699

**Authors:** Mahdi Dizani, Brian Perlstein, Diana McGrory, Syed Pavel Afrose, Siddharth Agarwal, Thomas Reese, Elisa Franco

## Abstract

Living cells rely on filaments and condensates as key organizers of their interior. Developing these structural primitives together inside synthetic compartments using programmable components is a step toward building functional synthetic cells, composite biomaterials, and synthetic tissues. Here, we demonstrate that DNA nanotubes and DNA condensates can be co-assembled within cell-sized compartments, including water-in-oil droplets and giant unilamellar vesicles (GUVs). The two nanostructures form as expected, producing a single condensate surrounded by nanotubes in diverse morphologies that depend on DNA and salt concentration, as well as compartment size. By incorporating photoactivatable DNA linkers, we can control the order of assembly and trigger reconfiguration of nanotube networks into ring-shaped bundles enclosing a condensate, architectures reminiscent of a cellular nucleus within a cytoskeletal ring. An isothermal assembly protocol based on monovalent salts further extends this approach to GUVs. Together, these results establish DNA filaments and condensates as programmable, composable organizers of synthetic cell interiors.

## Introduction

Numerous molecular compartments organize the interior of living cells. Cytoskeletal filaments partition cellular components, transport molecules, generate forces, and provide structural integrity^1–4^. Another important class of compartments is represented by biomolecular condensates, which have recently emerged as crucial regulators of diverse cellular reactions and processes^5–8^. Condensates typically present themselves as dense droplets that form via phase separation, and droplet nucleation, fusion, and dissolution occur dynamically in response to environmental stimuli^9,10^. Because filaments and condensates represent versatile “primitives” for cellular and biological organization, many synthetic platforms have been developed to characterize and engineer both^11–17^.

DNA nanotechnology provides versatile tools for designing and building molecular mimics of biological structures, including filaments and condensates^18,19^. Several works have demonstrated DNA-based nanotubes with physical properties and stiffness comparable to actin filaments^20–23^. Furthermore, the formation and dissolution of such nanotubes through physical and chemical stimuli have been demonstrated^24,25^. In parallel, DNA-based condensates have been built to exhibit liquid-like behavior similar to that of natural biomolecular condensates that undergo fusion, coalescence, and dissolution^26–30^. Synthetic DNA, or hybrid DNA-RNA condensates can also be used to recruit and localize proteins and target molecules, a function that is shared with some endogenous condensates^31,32^. Both classes of assemblies have been individually tested in water-in-oil droplets or vesicles to mimic biological compartmentalization^33–36^, paving the way to studies characterizing their co-assembly and mutual influence in confinement.

Here, we demonstrate that DNA filaments and condensates can be co-assembled both in bulk reactors and in microcompartments. Building on previous work, we produce nanotubes through the self-assembly of DAE-E tiles (i.e., nanotube monomers) consisting of five strands arranged into two parallel heteroduplexes connected by two crossover sites (Figure 1A, top). We produce DNA condensates using branched motifs known as DNA nanostars (i.e., condensate monomers), which form through the hybridization of three partially complementary strands (Figure 1A, bottom). Owing to the lack of base-pairing interactions between the two types of monomer, we find that the morphology of each type of structure is not affected by their co-assembly. Experiments in water-in-oil droplets reveal however that compartment size influences the morphology of DNA nanotube networks surrounding DNA condensates, across salt concentrations. We then introduce photoactivatable DNA linkers to generate ring-nucleus architectures. Our results are achieved by annealing monomers in divalent cation buffers, as well as through an isothermal protocol that relies on monovalent salts, which makes it possible to form nanotube-condensate systems in giant unilamellar vesicles (GUV). Collectively, our work shows that DNA filaments and condensates can be systematically combined as organizers of cell-sized compartments, and may be useful toward the development of synthetic cells.

**Figure 1.**
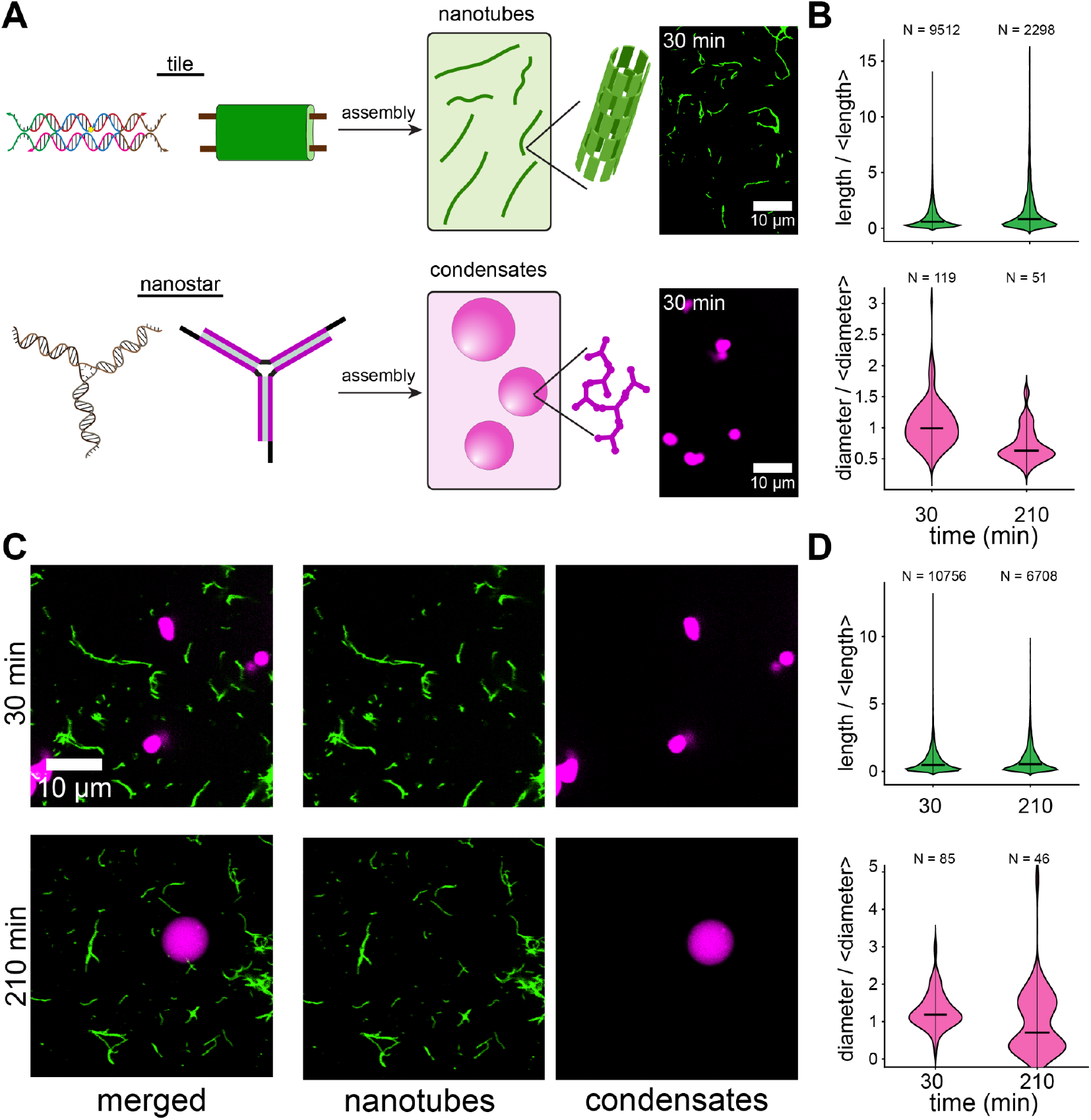
Co-assembly of DNA nanotubes and condensates in aqueous buffer. **A)** Schematic of DNA tiles and nanostars, which self-assemble and form nanotubes and condensates, respectively (left). Confocal microscopy images of nanotubes and condensates assembled separately (right). **B)** Violin plots depicting the distribution of nanotube length and condensate diameter normalized to the mean when assembled separately. **C)** Confocal microscopy images of nanotubes and condensates assembled together. **D)** Violin plots show similar size distribution of condensates and nanotubes when assembled together compared to when assembled separately. Experiments were run in duplicate (n = 2) and data was pooled and processed from both replicas.

## Results and Discussion

### Co-assembly of DNA Nanotubes and Condensates

We assembled together DNA nanotubes and condensates, adopting a 3-arm nanostar to form condensates^28,29^ and the DAE-E SEs tile to form nanotubes^20,37^ (Figure 1A). Although both motifs include multiple DNA strands (termed Y1, Y2, Y3 for nanostars, and S1 to S5 for tiles), they can be considered “monomers” that self-assemble into higher order structures via complementary domains, termed sticky ends. Nanostar sticky ends are single, palindromic domains, 4-nucleotides (nt) long (GCGC). DNA tiles present 4 sticky end domains, each 5-nt long, whose complementarity pattern leads to the assembly of curved lattices that fold into filaments, where DNA helices align with the filament principal axis. These sticky ends establish interactions between motifs of the same type, while interactions among nanostars and tiles are disfavored. We assembled both structures using the TrisHCl-TAE-MgCl2 buffer (the standard in the literature) and included distinct fluorescently labeled strands for sample imaging.

To control when condensate and nanotube assembly begins, we separately annealed incomplete monomers that cannot interact because they lack one of the sticky end carrying strands (Y2 for nanostars, S2 for tiles). Assembly is triggered when the missing strands are added to the sample containing incomplete monomers, and proceeds at room temperature. We assembled nanotubes and condensates both separately (Figure 1A, B) and together (Figure 1C, D), measuring their size 30 and 210 minutes after triggering their assembly. In the case where both nanostructures were present, no consistent interaction pattern was observed. Violin plots of nanotubes and condensates assembled together are comparable to those measured when the structures are formed separately. When both nanostructures are present, the persistence length of nanotubes measured from epifluorescence microscopy images on glass slides was 3.62 ± 0.15 µm and 3.54 ± 0.20 µm at 210 minutes for replicas 1 and 2, respectively (Figure S1, and Supplementary Methods). These measurements are consistent with the literature, indicating that the presence of condensates does not significantly affect the physical properties of nanotubes^20^.

### Growth of nanotubes and condensates in water-in-oil droplets

We next encapsulate tile and nanostar monomers in water-in-oil droplets^33,38^, with the goal of generating artificial structures that mimic nuclear compartments and cytoskeletal filaments inside biological cells. These droplets are prepared with a non-ionic surfactant and remain stable for weeks^39^. We produced droplets of uniform diameter of 17.50 ∓ 0.60 µm (averaged over 50 droplets) using a microfluidic chip (Figure 2A) and obtained a consistent morphology of nanotube “networks” surrounding a single DNA condensate, which resulted from the fusion of all smaller condensates emerging in the droplet^35^. When droplets were instead produced by vortexing solutions of nanostars and tiles together with an oil-surfactant mixture both the size of and the morphology within the emulsion droplets were polydisperse (Figure 2B)^38^. Because DNA nanostructure self-assembly depends on monomer abundance and ionic conditions, we varied the concentration of tiles (75, 150, and 300 nM), of nanostars (1.25, 5, and 15 µM), and MgCl_2_ (4 and 12.5 mM). Using the vortexing method (Figure 2B), we produced and then observed the encapsulated sample for up to 48 hours, and quantified the nanotube assembly by measuring the skewness of the distribution of pixel fluorescence intensity over time (in the nanotube channel).

**Figure 2.**
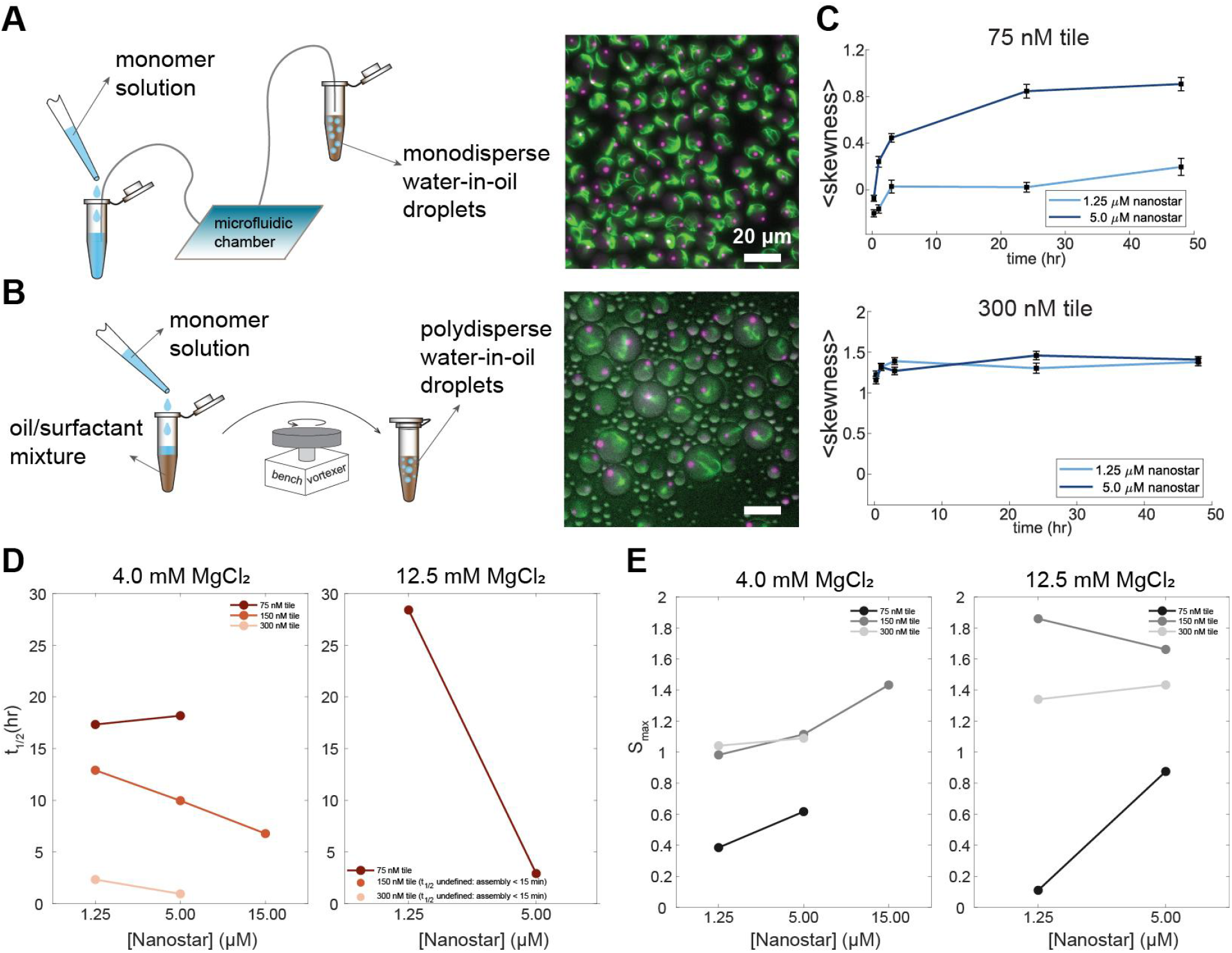
Isothermal self-assembly of nanotubes and condensates together in TrisHCl-TAE-MgCl_2_. **buffer A)** Encapsulation procedure of monomer solution using microfluidic chamber (left), and representative epifluorescence microscopy images of the monodispersed droplets 3 days after the sample preparation (right). **B)** Encapsulation procedure of monomer solution using a simple benchtop vortexer (left) and representative epifluorescence microscopy images of polydispersed droplets. **C)** Skewness of the histogram of pixel intensity of the nanotube channel, estimating the aggregation of nanotubes. Values averaged from n = 10 droplets with 8-12 μm diameter across 5 timepoints. Salt condition is at 12.5 mM MgCl_2_. The presence of nanostars accelerates aggregation at low tile concentration (75 nM tile) (top). At higher tile concentration, nanotubes aggregation is rapid and insensitive to nanostar level (bottom). **D)** Plots of the half-max time (t_1/2_ ) of the skewness time course (panel C) for tiles at different concentrations, under two salt conditions. **E)** Plateau skewness (S_max_) measured at the same conditions in D. See the Methods section for details about the plots.

The speed of nanotube assembly is primarily driven by the concentration of tiles and of MgCl_2_ . At low tile concentration (75 nM) and low MgCl_2_ (4 mM), nanotubes take more than 3 hours to form (Figure S2). Increasing tile and MgCl_2_ concentration accelerates nanotube growth (Figures S3, S4, and S5). Images suggest that higher nanostar concentrations promote nanotube assembly. This is confirmed by skewness measurements over time (Figure S6). This effect is more noticeable at low tile concentrations as compared to high tile concentration (Figure 2C), both in terms of the half-max time t_1/2_ and the steady-state skewness S_max_ (Supplementary methods), shown in Figure 2D and E. At 4mM MgCl_2_ and 75 nM tile level, S_max_ drastically increases with nanostar level while t_1/2_ remains comparable, suggesting assembly rates are unaffected even as the nanotubes themselves become more prone to aggregation. At 150 nM tile, t_1/2_ significantly decreases, while S_max_ remains comparable (Figure 2D, E and Figures S6 and S7).

### Morphology of nanotube networks in water-in-oil droplets

Following encapsulation and temperature reduction, condensates rapidly nucleate and fuse into a single spherical body within minutes. The size of the unified condensate depends on the nanostar concentration^35,40^. At the concentrations we explored, final condensate diameters were much smaller than the emulsion droplet diameter, which ranged from 5 µm to 20 μm. Thus, we expect that the size of the emulsion droplet has no major influence on the condensate morphology.

Unlike condensates, the morphology of nanotubes is influenced by the size of the emulsion droplet. We classified nanotube morphologies in three categories: rings, partial networks, and full networks. Because nanotubes move rapidly due to thermal fluctuations, we were not able to capture reliable confocal images, hence we report qualitative observations made from epifluorescence images. Rings are fluorescent objects consisting of one circular structure with no observed crossings between individual filaments. Partial networks have filaments that appear to intersect with at most 2 nodes. Full networks have more than 2 apparent filament intersections (Figure 3A and Figure S8). For each combination of MgCl_2_, tile, and nanostar concentration, 50 droplets were selected for morphology classification.

**Figure 3.**
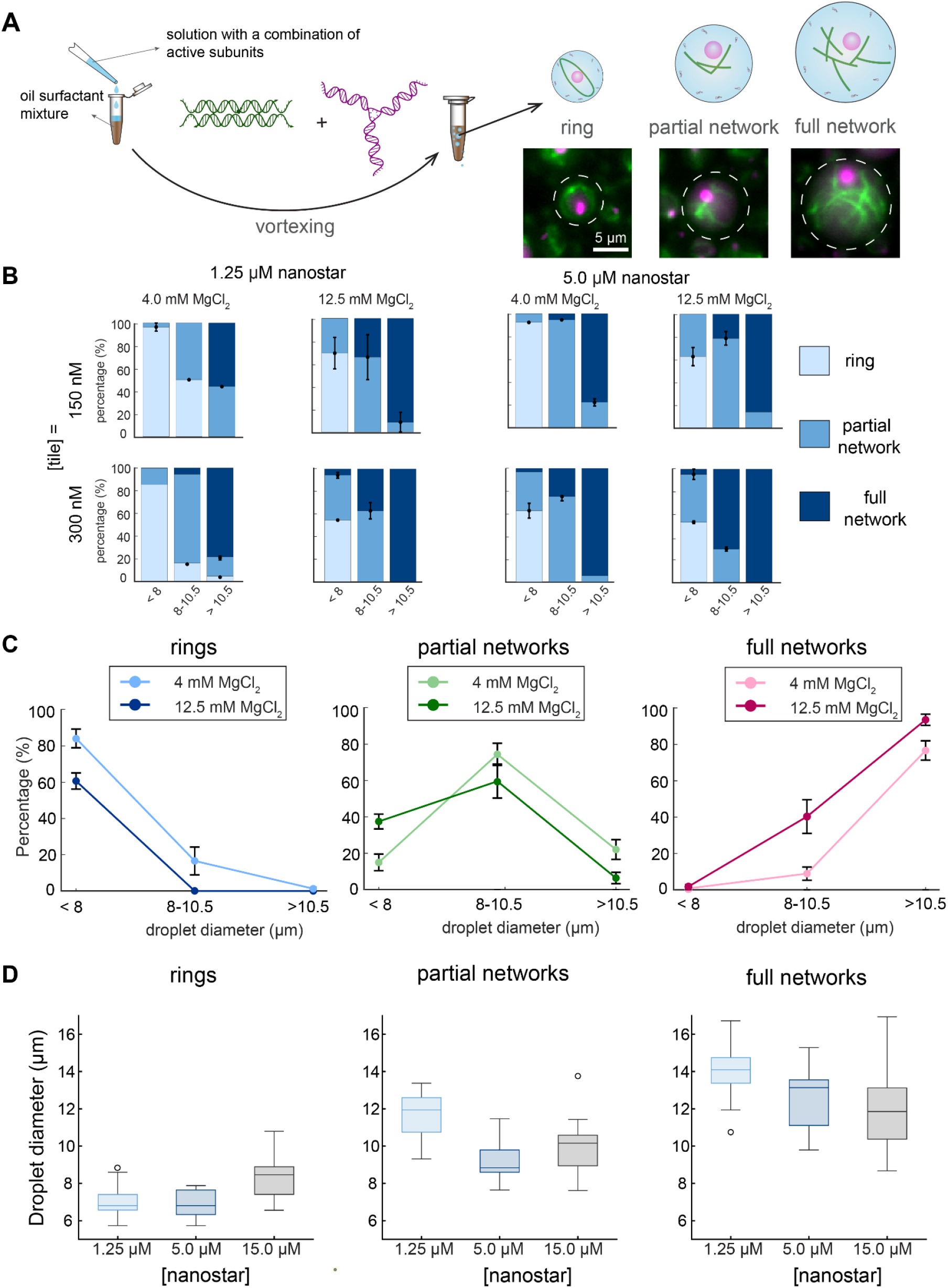
DNA scaffold morphology changes with droplet size and concentration of monomers and MgCl_2_. A) Schematic depicting how vortex emulsion results in polydispersal of droplet sizes and different nanotube morphologies in confinement. **B)** Bar plots demonstrating morphology categorization for nanotubes formed insides droplets binned in 3 diameter ranges (< 8, 8-10.5, and > 10.5 μm) for tile concentrations 150 nM (top) and 300 nM (bottom) and nanostar concentrations 1.25 μM (left) and 5 μM (right), comparing two different salt conditions, 4 and 12.5 mM. **C)** Plots showing the percentage of droplets that have a given nanotube morphology in each size range at 4 and 12.5 mM MgCl_2_. **D)** Box plots showing the distribution of droplet diameters corresponding to different nanotube morphologies, 150 nM tile and varying nanostar concentration. At 15 μM nanostar, rings form the most at the highest range of droplet diameter, while both partial and full network morphologies form at the highest range of droplet diameter at 1.25 μM nanostar.

Small emulsion droplets (< 8µm diameter) favor ring-like structures, whereas larger droplets (> 8 µm diameter) show more network formation (Figure S8). Increasing MgCl_2_ from 4 mM to 12.5 mM shifts the morphology toward partial and full networks across all droplet sizes while reducing the occurrence of the ring morphology (Figure 3B, C). The formation of rings in small droplets is likely because the persistence length of nanotubes (3.62 and 3.54 µm) has a similar scale as the droplet diameter, which results in bending and alignment of nanotubes. Increasing the salt concentration likely facilitates nucleation, leading to shorter nanotubes, which adopt a network-like configuration. Nanostar concentration also influences nanotube morphology. At 1.25 µM nanostar concentration networks prevailed, while at 15 µM nanostars ring morphologies occur in relatively large emulsion droplets (Figure 3D). It is known that molecular crowders induce ring formation through depletion interactions^41^, and our results indicate this effect depends on the crowder concentration. Increasing nanostar concentration also facilitates the emergence of networks, but Interestingly, at 15 μM nanostar, ring morphologies occur in relatively large emulsion droplets, whereas networks prevail at 1.25 μM nanostar concentration (Figure 3D).

### Timing the assembly of nanotubes and condensate through UV irradiation

In biological cells, the assembly of condensates and cytoskeletal filaments can be independently regulated in response to distinct stimuli, motivating the design of synthetic systems where each structural phase can be triggered separately. We achieved this by modifying DNA tiles and nanostars to include domains that respond to ultraviolet (UV) light irradiation, building on previous work^25,35^. We included a photocleavable (PC) hairpin domain on the S2 strand of tiles and on the Y2 strand of nanostars (Figure 4A)^35^. By occluding one sticky end, the PC hairpin produces a “caged” motif with reduced valency that cannot self-assemble. UV irradiation causes the PC hairpin to cleave and dissociate from the sticky end, which becomes available for binding other sticky ends. Irradiation effectively increases the valency of the motif, enabling assembly of the expected nanostructures (Figure 4A).

**Figure 4.**
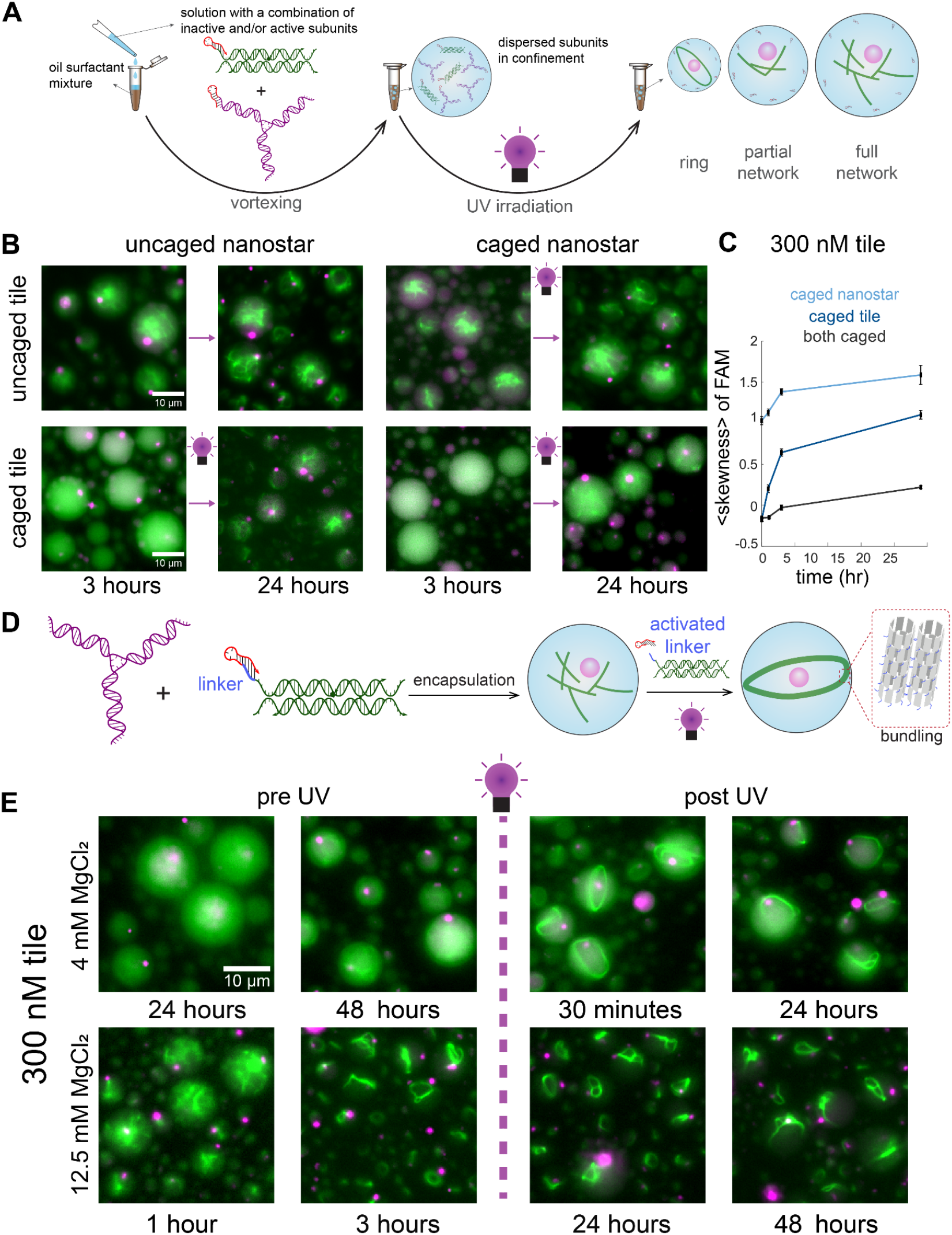
Photoactivatable DNA tile and nanostar motifs display latent formation after light irradiation and can be used to induce morphological changes in confinement. **A)** Addition of a photocleavable domain on one sticky end of tile strand S2 and nanostar Y2 prevents formation of either structure until irradiation of UV light. FAM channel of nanotubes shown in green and Cy5 channel of condensates shown in magenta. **B)** Epifluorescence microscopy images demonstrate condensate formation (magenta) and nanotube growth (green) upon UV irradiation. Images were taken at 300 nM tile and 4 mM MgCl_2_. **C)** Skewness of the pixel intensity histogram^33^ is used as an estimate for the aggregation of nanotube networks. Values were averaged for 10 droplets from 8-12 μm in diameter across 5 timepoints. Images were taken 3 hours after forming emulsion droplets and before irradiation, indicating timepoints as time after UV irradiation. **D)** Addition of a linker toehold at the end of the sticky end domain allows for bundling and formation of rings in nanotube networks. A photocleavable hairpin occludes the linker to induce bundling of nanotube networks upon UV irradiation. **E)** At 300 nM tile and 4 mM MgCl_2_, nanotube networks form after 48 hours and irradiation demonstrates bundling of nanotubes into rings. At 12.5 mM MgCl_2_, full networks form after 3 hours with 150 nM and 300 nM tile concentrations. Nanotubes bundle 3 hours post-irradiation and retain a ring-like morphology.

By making emulsion droplets that include different combinations of caged motifs, we can produce samples in which UV irradiation triggers the formation of nanotubes, condensates, or both (Figure 4B). In these experiments, the uncaged motif begins assembling as soon as the sample is prepared, while the caged one does not unless UV irradiation is provided. This allows for selectively producing an artificial “nucleus” or “cytoskeleton” in the emulsion droplets only after the UV stimulus is applied. We characterized the assembly process at different tile and nanostar concentrations (Figures S9-S12).

UV-activated tiles assemble less efficiently in samples where tiles are caged while UV-activated nanostars robustly form condensates (Figure 4C). We hypothesize that dissociated PC hairpin weakly interacts with the 5 nt-long sticky ends on tiles and thereby reduce the tile assembly rate, limiting the speed of nanotube formation, but have negligible interaction with the 4nt-long sticky ends on nanostars. Interestingly, nanotube assembly after UV irradiation appears least efficient when both tiles and nanostars are caged (Figures 4C, S13), an effect that persists up to 24 hours (Figure 4B, 4C). As expected, increasing tile concentration promotes the speed of nanotube formation, although very high tile concentration (300 nM) generates significant aggregation and few discernible nanotubes (Figures S12).

### Photo-controlled nanotube bundling yields ring-nucleus morphologies

To mimic the capacity of cells to remodel their cytoskeleton in response to stimuli, we modified DNA tiles to include a photocleavable palindromic domain on the 5’ end of strand S2, exposing a linker on the outer nanotube surface upon UV irradiation^42^. This strategy makes it possible to use UV irradiation to rearrange nanotube networks into bundles that wrap into a single ring that usually adsorbs on the emulsion droplet surface (Figures 4E, S14). The presence of uncaged nanostars consistently yields a single condensate, reminiscent of a nucleus, complemented by a thick nanotube ring that we term “Saturn” morphology. Because bundling is irreversible in our design, droplets retain the Saturn morphology long-term (measured up to 48 hours, Figure S15). Increasing the concentration of MgCl_2_ accelerates the assembly and aggregation of nanotubes into full networks, and this conformation hinders ring formation after UV irradiation (Figure 4E, bottom) when compared to a low MgCl_2_ condition.

### Isothermal co-assembly of nanotubes and condensates in vesicles

Finally, we demonstrate the co-assembly of nanotubes and condensates at constant room temperature in Giant Unilamellar Vesicles (GUVs). To this end, we adopted the TANa buffer^43^ for isothermal assembly (40 mM Tris HCl, 20 mM acetic acid, and 100 mM NaCl), and confirmed that nanotubes and condensates both assemble in this buffer (Figure 5A) on a timescale comparable to that observed under anneal-conditions with TrisHCl-TAE-MgCl_2_ buffer (Figure 1A). Nanotube length violin plots show consistent distributions over time, however condensate diameter distributions at 210 minutes deviate significantly from the 30 minute timepoint (Figure 5B). In TANa buffer, the persistence length of nanotubes at 210 minutes was 7.76 ± 0.89 µm and 9.09 ± 1.01 µm for replicas 1 and 2, respectively, notably higher than values measured in MgCl_2_ buffer. As a control, we then verified that TANa buffer allows for nanotube and condensate co-assembly in water-in-oil droplets. As expected, nanotube assembly is faster and more networks are observed when operating at high tile concentration (Figure 5C), while low tile concentration (75 nM) results in little growth even after 48 hours (Figures S16-S17).

**Figure 5.**
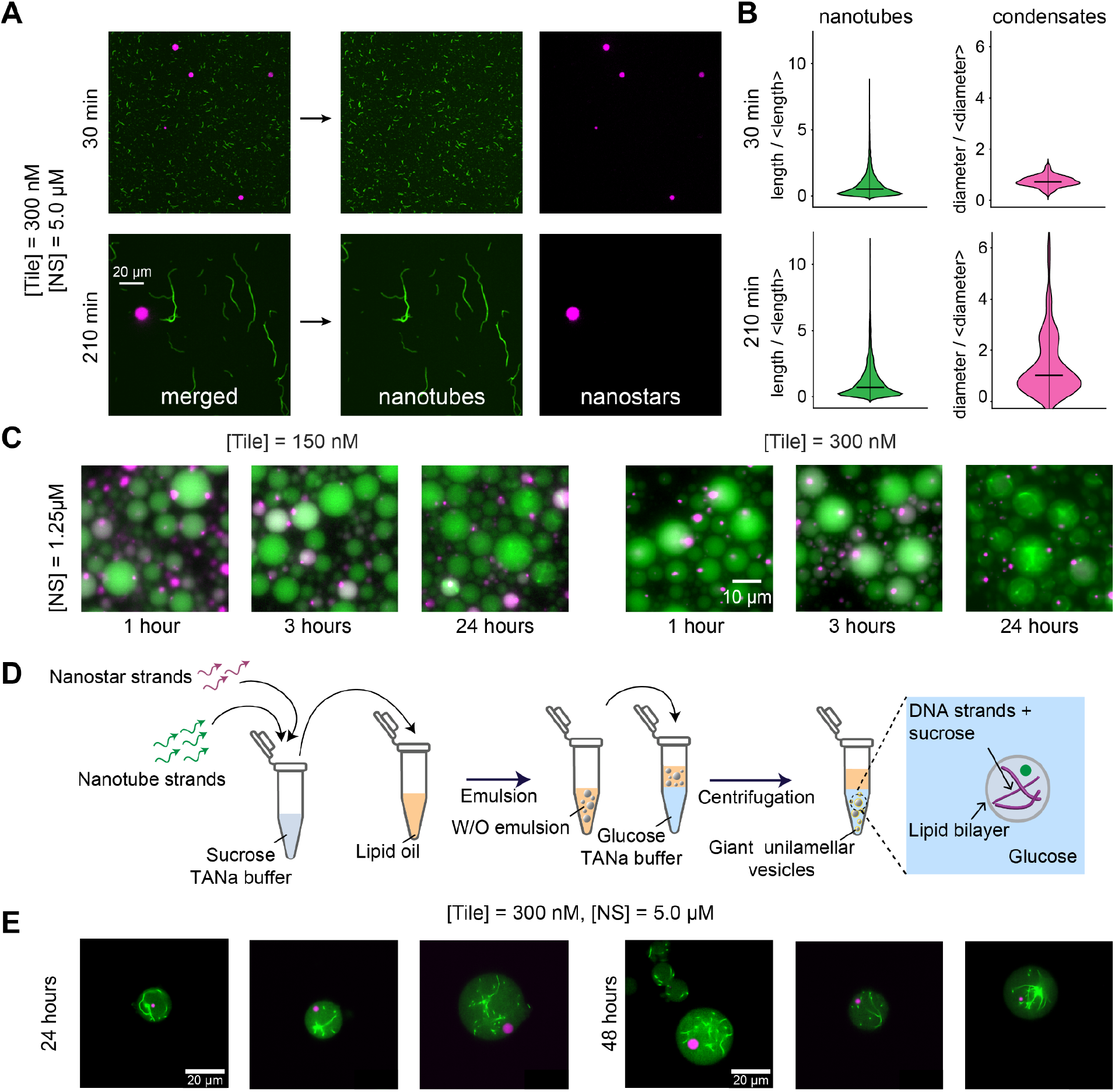
Isothermal growth of nanotubes and condensates in w/o droplets and giant unilamellar vesicles (GUVs) using TANa buffer. **A)** Confocal microscopy images depicting isothermal formation and growth of DNA condensates and nanotubes in TANa buffer at 200 mM NaCl at 30 minutes and 210 minutes post mixing. **B)** Violin plots of the size distribution of condensates and nanotubes at different time points. **C)** Epifluorescence microscopy images showing the isothermal growth of nanotubes and condensates in water-in-oil droplets at [tile] = 150 nM (left) and 300 nM (right). **D)** Schematic of the formation of GUVs for the encapsulation of nanotubes and condensates. Emulsion and centrifugation produces bilayered vesicles incorporating nanostar and tile monomers for isothermal growth. **E)** At 24 and 48 hours, nanostars phase separate into condensates and nanotubes begin growing into networks.

Nanotubes and condensate assemble as anticipated in GUVs produced through the double-emulsion method^43,44^ (Figure 5D), which generates a polydisperse vesicle population with equivalent diameters ranging from approximately 4 to 74 µm, with most vesicles exhibiting diameters below ∼20 µm. Across all vesicle diameters, condensates and nanotubes formed at 24 hours, retaining their morphology up to 48 hours (Figure 5E). At 1µM tile concentration, nanotube networks appear more abundant when compared to a 300 nM tile condition. In some cases, a high concentration of tiles results in nanotube aggregation (Figure S18).

## Conclusions

We demonstrated that DNA nanotubes and condensates can be co-assembled inside cell-sized compartments as independently programmable structural phases. The orthogonality of their sticky end interactions ensures that the two assemblies coexist without molecular cross-talk, retaining size statistics comparable to those they exhibit when assembled in isolation. When encapsulated in emulsion droplets, condensates reliably fuse into a single body while nanotubes adopt morphologies (rings or networks) governed by compartment size, tile concentration, and salt content. By incorporating photocleavable hairpins, we demonstrate on-demand triggering of either phase independently, and the sequential assembly of nanotube networks followed by UV-induced bundling to consistently produce a “Saturn” morphology: a single condensate and a nanotube ring. Together these results establish a modular toolkit for programming the internal architecture of synthetic cells, where the two structural phases can be designed, triggered, and reconfigured.

A notable finding is that nanostar concentration affects nanotube assembly, particularly at low tile concentrations. Quantifying nanotube assembly (through the half-max time t_1/2_ and the plateau S_max_ of pixel intensity skewness) across conditions reveals that increasing nanostar concentration enhances both the speed and the extent of assembly at low tile concentration. Several mechanisms could explain these observations. Excluded volume effects induced by nanostars could raise the effective local tile concentration and lower the nucleation barrier, a well-documented phenomenon in crowded macromolecular environments. Alternatively, nanostars could alter tile-tile interactions through changes in the local electrostatic environment, or promote weak nonspecific tile clustering that lowers the nucleation barrier. Among these, excluded volume is arguably the most parsimonious explanation, as the nanostar effect is strongest at low tile concentrations where the nucleation barrier is highest and most sensitive to changes in effective monomer concentration. Distinguishing between these mechanisms remains an open question.

Compared to prior studies that encapsulated DNA nanotubes or condensates individually in compartments, our work shows that the two phases can be combined and that their co-existence is governed primarily by physical confinement rather than molecular cross-talk. Previous work showed assembly of a DNA cytoskeleton in confinement^33,34,43^, or formation of DNA or RNA condensates in confinement^35,45,46^, but neither combined both structural primitives nor characterized their mutual influence. The extension to GUVs via the TANa isothermal protocol is practically important, as it makes the system compatible with lipid membrane compartments that are sensitive to divalent cation conditions. The use of UV irradiation to control the sequential assembly and morphological reconfiguration of internal compartment architecture represents a particularly timely contribution. Light has recently been highlighted as a uniquely precise, biorthogonal, and non-invasive stimulus for regulating synthetic cell systems, including compartmentalization, shape change, and the establishment of spatial and temporal inhomogeneity^47,48^. Our results extend these advances in two complementary directions. First, they demonstrate selective on-demand triggering of a single structural phase while leaving the other unperturbed, a degree of orthogonal control that is difficult to achieve with chemical or thermal stimuli. Second, they demonstrated a morphological reconfiguration, switching filaments into bundles. This capacity to not only trigger but also reconfigure the internal architecture of a compartment after assembly is a key step toward synthetic cells that can dynamically reorganize their internal structure in response to external cues, as living cells do. A current limitation is the irreversibility of the bundling step in our design; future work incorporating reversible photoswitches such as azobenzenes could enable repeated cycles of network formation and ring reconfiguration^49^. More broadly, the modularity and programmability of the DNA-based platform established here opens the door to incorporating biochemical functionality^31,50^ (for instance by recruiting proteins or nucleic acid circuits to the condensate phase) and to studying how spatial organization of internal compartments influences the reactions they contain, questions that are difficult to dissect in natural cells but tractable in this bottom-up setting.

## Methods

### Oligonucleotides

DNA strands (standard desalted) were purchased from Integrated DNA Technologies IDT, Coralville, IA, USA. Fluorescently labeled strands were HPLC-purified. Nanostar strands are standard desalting and tile strands are PAGE-purified. Oligonucleotides were quantified using a Thermo Fisher Scientific NanoDrop 2000 Spectrophotometer. Strands were suspended in nanopure water and stored at -20 °C for long-term usage. Sequences were listed in the Supporting Information (Table S1).

### Nanotube and condensate preparation

The core strands of the nanostars (Y1 and Y3) were annealed in 20 mM Tris-HCl, 3 mM MgCl_2_ then kept at room temperature for isothermal addition of strand Y2. Annealing was performed using a Bio-Rad T100™ Thermal Cycler, ramping up to 90°C, then cooling down by -1 °C/min to 25°C. In order to anneal tiles, S1, S3, S4, and S5 were annealed in 5 mM MgCl_2_ and 1X TAE and then stored at room temperature. Tiles annealing was done by heating to 90 °C, then cooling to 25°C over 6 hours. Nanostars and tiles were mixed at appropriate concentrations in the mixture of TrisHCl-MgCl_2_ (nanostars buffer) and TAE-MgCl_2_ (tiles buffer) were adjusted to obtain a final composition of 1X TAE, 20 mM Tris-HCl, and different MgCl_2_ levels. All experiments were completed in duplicate (n = 2) unless otherwise noted. For TANa buffer experiments, we used 40 mM Trizma-base, 20 mM acetic acid, and 100 or 200 mM NaCl. In experiments employing the photocleavable hairpin, the sample was irradiated for 180 s approximately 4 cm away from a sealed Ibidi chamber (μ-Slide VI 0.4) by a 302 nm, 8 Watt light source with 115 V, 60 Hz (UVP 3UV Lamp, Analytik Jena).

### Preparation of water-in-oil droplets

The aqueous phase including nanostar and tiles at desired concentrations and buffer composition was heated to 55°C and mixed with a micropipette prior to encapsulation to ensure full dispersion of monomers. Then, 20 µL of each sample was added into 0.5 mL Eppendorf LoBind® tubes directly on top of 80 µL of 2% weight per volume oil-surfactant mixture. The oil-surfactant was made with a 2% (w/v) mixture of oil (FC-40 Fluorinert™, Sigma-Aldrich) and surfactant (008 FluoroSurfactant, RAN Biotechnologies). The sample was then vortexed for 44-48 seconds, then equilibrated for the same time period to generate droplets with a polydispersed size range. A 50 µL aliquot of the milky fraction of the sample was transferred into an Ibidi chamber (μ-Slide VI 0.4) then sealed with Molykote vacuum grease and covered with a coverslip to prevent evaporation of the samples.

To generate monodispersed droplets, we employed a microfluidics setup (LabSmith, Inc.). Droplets were formed by interfacing a pressurized continuous (oil-surfactant) phase and dispersed (DNA-laden aqueous solution) phase to meet within a commercial droplet generating chip of a 20 micron diameter intersection (Chipshop, Fluidic 947). The microfluidics stage was heated up to 50 °C to ensure that the monomers remain dispersed in the solution and do not self-assemble. The emulsion mix was then collected in the test tube and was transferred to chambers, sealed with grease for long-term imaging.

### Preparation of lipid vesicles

A chloroform solution of DOPC was placed in a glass vial and the chloroform was removed by blowing nitrogen for 5 minutes. The film was resuspended in mineral oil (M5904, Sigma) by sonication for 30 min, to a final concentration of 1.33 mg/mL. The composition of the vesicle outside solution was 40 mM Trizma base, 20 mM acetic acid, 100 mM NaCl and 232.5 mM glucose. In the inside vesicle solution, glucose was replaced by 230.1 mM sucrose, with the addition of DNA strands at the desired concentrations. The inside sucrose solution was first emulsified by pipetting in the lipid oil solution, to a ratio of 1:30, followed by 1 minute of vortexing. The emulsion was slowly added in a lipid oil solution placed on top of the outside glucose solution. These water-in-oil droplets were left sedimented for 5 minutes at the water-oil interface, before being centrifuged at 4500 rcf for 5 minutes at 5 °C. The oil solution was removed and the outside solution containing the vesicles was imaged. Vesicle imaging was performed in untreated 8-well chambered coverslips (ibidi μ-Slide 8 Well High; polymer coverslip, 180 ∓ 10.5 μm). Before imaging, the wells were incubated with 3% (w/v) BSA to passivate the surface and reduce vesicle adhesion and washed with the glucose containing buffer (the outside solution).

### Microscopy imaging

For the aqueous buffer experiments, samples were imaged with a Nikon Eclipse Ti2, using a Nikon CFI Plan Apo Lambda 60X Oil objective, and an Eclipse filter cube with excitation wavelength of 448 nm for FAM-labeled tiles and 646 nm for Atto 647-labeled nanostars. Images of the GUVs were acquired using a Nikon Ti confocal microscope equipped with an NL5+ camera and a 60X oil-immersion objective. Before imaging the bulk samples, coverslips (Fisherbrand™, 60×22 mm, 0.13 to 0.17 mm thick) were utilized to create imaging chambers, prepared with a Parafilm (Fisher Scientific) squared piece with a punched hole on it, adhered to the coverslips by heating to 50-60°C. After the coverslip cooled down, 2.5 μL of the sample was loaded in the punched region. To minimize evaporation, the sample was then covered with a smaller coverslip (Fisherbrand™, 22×22 mm, 0.13-0.17 mm thick).

### Image analysis

For experiments in aqueous buffer, six microscopy images were taken from each replica and were processed using in-house code to detect and measure the length scale of both nanotubes and condensates. The code is available on GitHub at: https://github.com/klockemel/. Skewness measurements were obtained via Fiji ImageJ by selecting 10 circular ROIs based on the droplet size across two replicas for each timepoint. Large droplets (> 10.5 µm) of a similar size across all timepoints in a given experiment were considered. The computation of half-max time t_1/2_ and plateau skewness S_max_ is described in the Supplementary methods.

Nanotube persistence length was measured as described in the Supplementary methods. Nanotube morphology in confinement was examined using images taken of emulsion droplets incubated for 1 day. A total of 50 droplets were selected from different experimental replicas, choosing 15-20 droplets in each diameter category and manually identifying the nanotube network morphologies (Figure S8). All the processed data were plotted using MATLAB.

## Associated Content

## Supporting Information

Supplementary tables; supplementary methods; supplementary figures.

## Acknowledgements

We thank Deborah K. Fygenson for helpful discussions and advice. This research was supported by the Department of Energy (DOE), Office of Science, Basic Energy Sciences (BES), Award SC0010595 to EF and by the Sloan Foundation through award G-2024-22575.

The authors used Claude Sonnet 4.6 and ChatGPT for editing the text and for drafting some of the data processing code. After using these tools, the authors reviewed and edited the content as needed and take full responsibility for the final content of this manuscript.

