## Supplementary tables; supplementary methods; supplementary figures. for "Programming the Internal Architecture of Synthetic Compartments by Coassembling Filamentous and Liquid DNA Phases"

**1. Supplementary tables** **2**

**2. Supplementary methods** **3**

**3. Supplementary figures** **5**

**4. References** **22**

### 1. Supplementary tables

**Table S1. Oligonucleotides used in the study.** Sequences used to form tiles and nanostars were adapted from our previous studies[1,2].

| Name (Tile) | 5' – sequence – 3' |
|---|---|
| **S1 (5bSE1)** | CTCAGTGGACAGCCGTTCTGGAGCGTTGGACGAAACT |
| **S2 (5bSE2)** | GTCTGGTAGAGCACCACTGAGAGGTA |
| **$S2_{photo}$ (5bSE2_Photo)** | GTCTGGTAGAGCACCACTGAGAGGTA/iSpPC/GGTAAAACCTAC |
| **$S2_{linker}$** | **CGCG**TTTGTCTGGTAGAGCACCACTGAGAGGTA |
| **$S2_{linker,photo}$** | GCGCCAAAATGG/iSpPC/**CGCG**TTTGTCTGGTAGAGCACCACTGAGAGGTA |
| **S3 (5bSE3_fam)** | /**fam**/CCAGAACGGCTGTGGCTAAACAGTAACCGAAGCACCAACGCT |
| **S4 (5bSE4)** | CAGACAGTTTCGTGGTCATCGTACCT |
| **S5 (5bSE5)** | CGATGACCTGCTTCGGTTACTGTTTAGCCTGCTCTAC |
| **Name (NS)** | **5' – sequence – 3'** |
| **$Y1^{atto647}$** | /**atto647**/CAGTGAGGACGGAAGT**TT**GTCGTAGCATCGCACC |
| **Y1** | **GCGC**CAGTGAGGACGGAAGT**TT**GTCGTAGCATCGCACC |
| **Y2** | **GCGC**CAACCACGCCTGTCCA**TT**ACTTCCGTCCTCACTG |
| **$Y2_{photo}$** | CGCACCAAAGGT/iSpPC/**GCGC**CAACCACGCCTGTCCA**TT**ACTTCCGTCCTCACTG |
| **Y3** | **GCGC**GGTGCGATGCTACGAC**TTT**GGACAGGCGTGGTTG |

### 2. Supplementary methods

#### Persistence length estimation of nanotubes

Persistence length was calculated using epifluorescence microscopy images of DNA nanotubes deposited on glass slides and the images were analyzed using a custom Python pipeline to extract filament contours and quantify persistence length (Lp). For samples at a specific time point, 10 independent fields of view (FOVs) were analyzed.

Standard scientific Python libraries  such as NumPy, SciPy, scikit-image, and NetworkX were used for the image processing and analysis steps. Raw images were converted to floating-point intensity arrays and were normalized by their maximum intensity. Images were smoothed using a Gaussian filter ($\sigma$ = 1 pixel) to suppress high-frequency noise. Background signal was estimated using morphological opening with a disk structuring element (radius = 15 pixels) and subtracted using a scaling factor of 0.7 to enhance filament contrast. Binary masks of filamentous structures were generated using hysteresis thresholding, with lower and upper thresholds set to 0.2 and 0.5 times the maximum image intensity, respectively. To suppress noise and debris, small connected components containing fewer than 32 pixels (corresponding to an area of approximately 0.16 $\mu m^2$) were removed. Binary masks were skeletonized to produce one-pixel-wide centerlines of filament structures.

Skeletons were screened for branch points using morphological hit-or-miss operations with multiple rotated structuring elements to ensure analysis of individual, non-intersecting filaments. Any connected component containing one or more branch points (i.e., intersections or junctions) was discarded. This step was performed to ensure that only isolated, linear nanotube segments were retained for analysis.

Each connected skeleton component was converted into a graph representation, where pixels were treated as nodes and edges were assigned Euclidean weights based on pixel connectivity. Candidate paths between endpoints were identified, and the longest geodesic path within each component was selected as the representative filament backbone. The selected path was then converted into a continuous polyline in physical units using a calibrated pixel size of 0.07 μm $pixel^{-1}$. Consecutive duplicate coordinates were removed prior to calculation of cumulative contour distance. The cumulative arc length along this polyline yielded the contour length of each filament. Only filaments exceeding a minimum contour length threshold (≥0.3 μm) were included in further analysis.

Local tangent vectors along each filament were computed using finite differences of the polyline coordinates, followed by normalization to unit vectors. The tangent–tangent correlation function was then calculated as a function of contour distance:

$$C(\Delta s) = \langle t(s) \cdot t(s + \Delta s) \rangle$$

where t(s) is the unit tangent vector at contour position s. Correlations were computed by averaging dot products of tangent vector pairs separated by a given arc length. The correlation function was evaluated up to 50% of the total contour length to minimize finite-size effects and was binned uniformly along the contour into 60 contour-distance bins.

Persistence length (Lp) was determined by fitting the tangent–tangent correlation function to the worm-like chain (WLC) model:

$$C(\Delta s) = \exp(-\Delta s/Lp)$$

Fitting was performed using nonlinear least-squares regression. Only finite correlation values satisfying $0 < C(\Delta s) \leq 1$ were included in the fitting procedure, and fits were performed only when at least five valid correlation points were available. Initial parameter estimates were obtained from a linear fit to the logarithm of the correlation function. The quality of each fit was quantified using the coefficient of determination ($R^2$).

Representative tangent–tangent correlation plots shown in Figure S1 were generated by pooling nanotube correlation data across all nanotubes from the analyzed fields of view within an experimental replicate and are provided to visualize the overall correlation decay and fitting quality. These ensemble-averaged fits were used only for illustrative purposes and were not used for the quantitative determination of sample-level persistence lengths. Persistence lengths reported in this work were instead obtained by fitting individual nanotubes, calculating the median individual-filament Lp within each field of view, and subsequently reporting the mean ± standard error of the mean (SEM) across the independent fields of view.

**Quantification of Nanotube Assembly Kinetics**

Nanotube assembly kinetics were estimated by measuring the skewness of the fluorescence intensity histograms measured across droplet regions of interest at five timepoints (15, 60, 180, 1440, and 2880 minutes). Skewness was measured for n=10 droplets per condition and timepoint, and the mean skewness was computed across droplets at each timepoint.

Two summary statistics were extracted empirically from the mean skewness curve. The plateau skewness $S_{max}$ was defined as the mean of the last two timepoints (1440 and 2880 minutes), providing an estimate of the maximum degree of nanotube network formation achieved under each condition. The assembly half-time $t_{1/2}$ was defined as the time at which the mean skewness curve crosses $S_{max}/2$, estimated by linear interpolation between the two bracketing timepoints. For conditions where the skewness already exceeded $S_{max}/2$ at the first timepoint (15 minutes), $t_{1/2}$ was considered undefined, indicating that assembly was essentially complete before the first observation.

This empirical approach was preferred over parametric alternatives such as sigmoidal or exponential fitting for two reasons. First, with only five timepoints spanning three orders of magnitude in time, the data are too sparse to reliably identify the parameters of a kinetic model, and fitted parameters carry large uncertainties that are not physically meaningful. Second, the empirical estimates make no assumptions about the functional form of the assembly kinetics, making them agnostic to mechanistic differences across conditions.

The resulting plots of $t_{1/2}$ and $S_{max}$ versus nanostar concentration allow two distinct aspects of assembly to be compared across conditions: $t_{1/2}$ reports on the speed of assembly onset, revealing how tile, nanostar, and salt concentration influence the kinetics of network formation. $S_{max}$ reports on the extent of assembly at steady state, reflecting whether the final network morphology depends on the same parameters. Together, these metrics provide a compact and interpretable summary of how environmental conditions govern both the dynamics and the outcome of nanotube assembly inside synthetic compartments.

#### 3. Supplementary figures

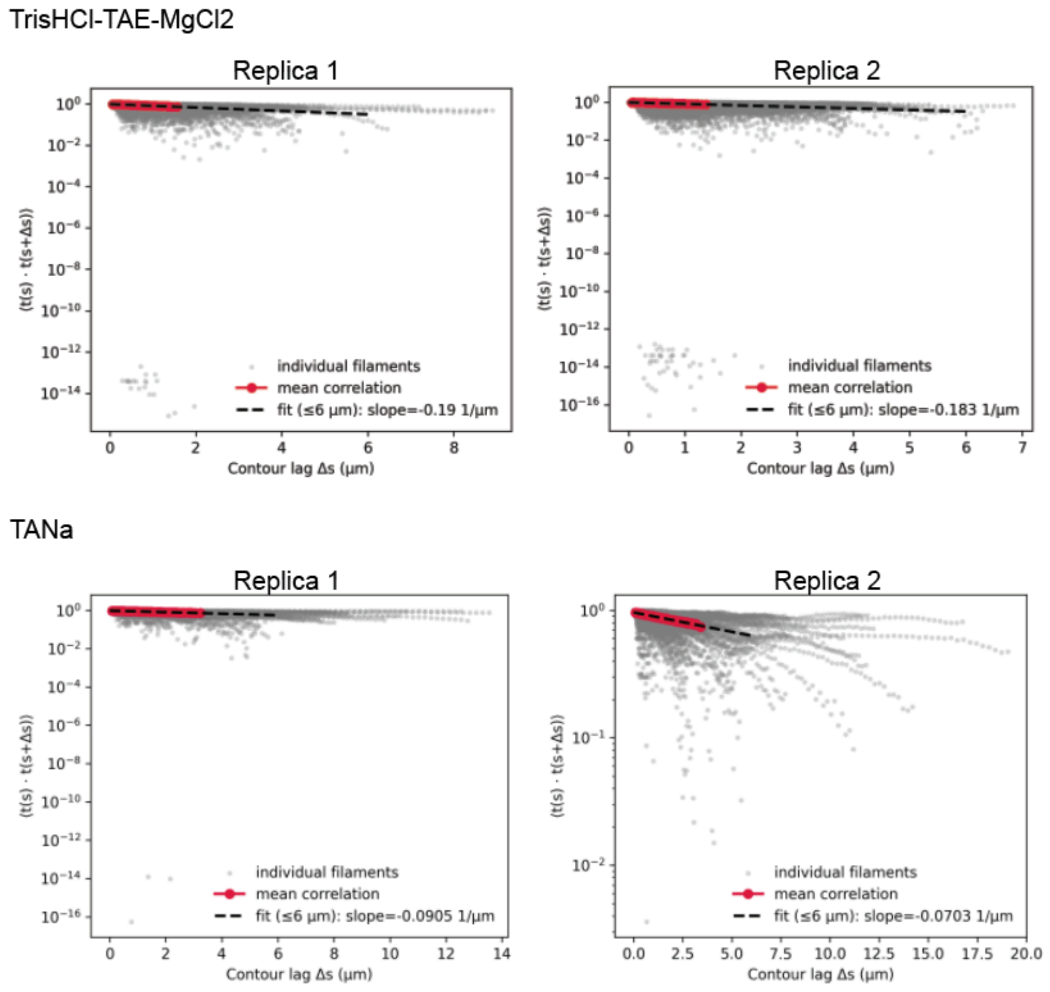


***Figure S1. Ensemble tangent–tangent correlation analysis for independent experimental replicates in different buffer conditions.*** *For each replicate, nanotube contours extracted from 10 independent fields of view were pooled prior to correlation analysis. Gray points represent tangent-correlation values from individual nanotubes, whereas red points indicate the mean correlation at each contour-distance bin across all nanotubes within a replicate. Black dashed lines show linear fits to ln C(Δs) over 0 < Δs ≤ 6 μm, with the fitted slopes indicated. The corresponding apparent ensemble persistence length is given by Lp = −1/slope. These ensemble-averaged plots are shown solely to illustrate the overall decay of tangent correlations and fitting quality. Persistence lengths reported in the manuscript were instead obtained by fitting individual nanotubes, calculating the median individual-filament Lp for each field of view, and subsequently reporting the mean ± SEM across the 10 independent fields of view within each replicate.*

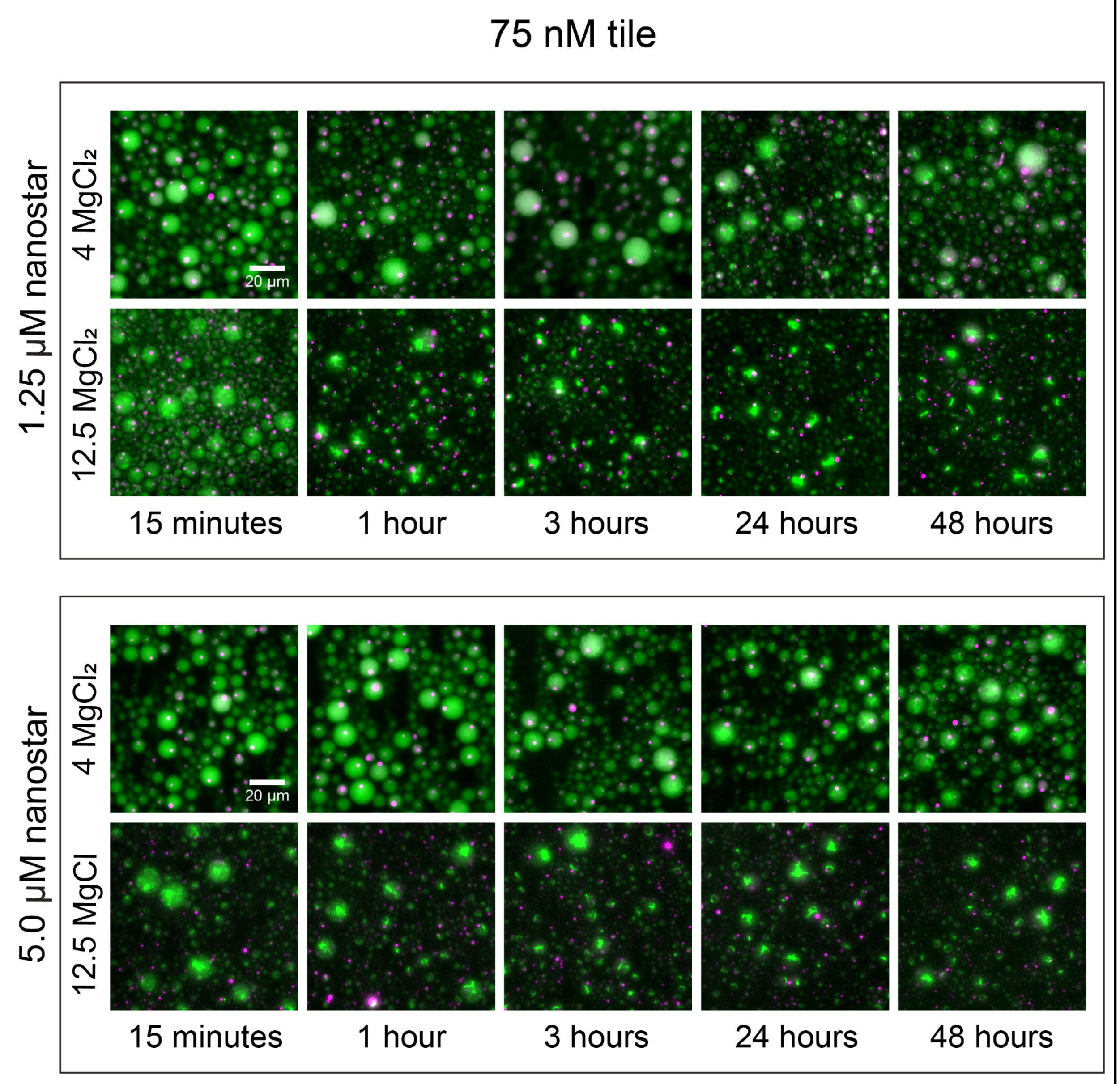


***Figure S2. Kinetic images of water-in-oil droplets show growth of condensates and nanotubes at 75 nM tile.*** *At a low concentration of tile, there is significant variability in nanotube growth depending on nanostar and* $MgCl_2$ *concentration. Increasing nanostar concentration promotes faster and more robust assembly of nanotubes. Increasing* $MgCl_2$ *concentration has a more significant effect on the nanotube self-assembly rate.*

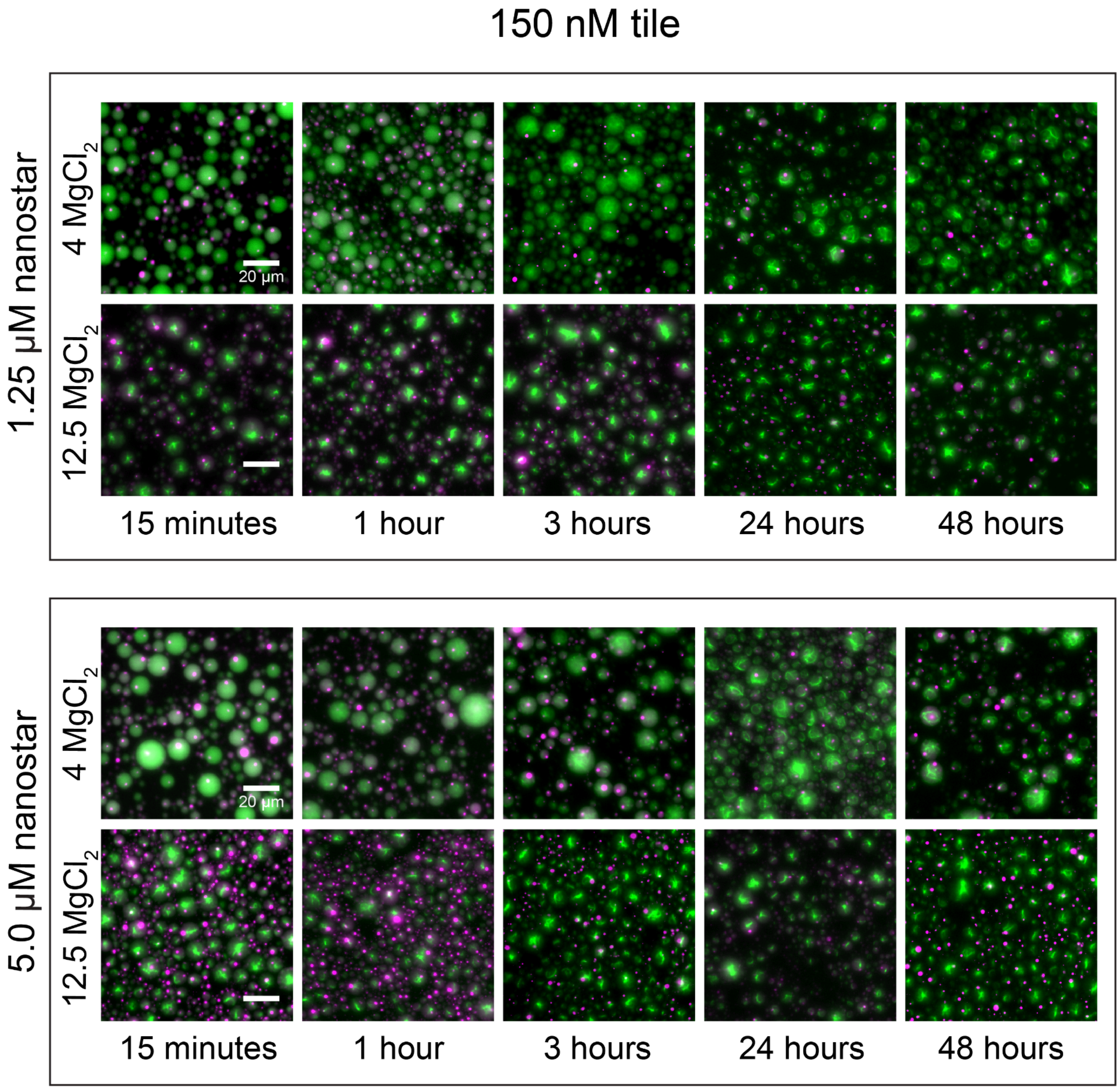


***Figure S3. Kinetic images of water-in-oil droplets show growth of condensates and nanotubes at 150 nM tile depending on different nanostar and $MgCl_2$ concentrations.*** *Increased concentration of $MgCl_2$ from 4 to 12.5 mM facilitates initial nucleation of tiles and rapid growth of nanotubes in water-in-oil droplets. At higher $MgCl_2$, nanotubes form more aggregates, rather than clean networks found at lower $MgCl_2$. Condensates form rapidly within 15 minutes of water-in-oil droplet emulsion and retain their size up to 2 days after formation.*

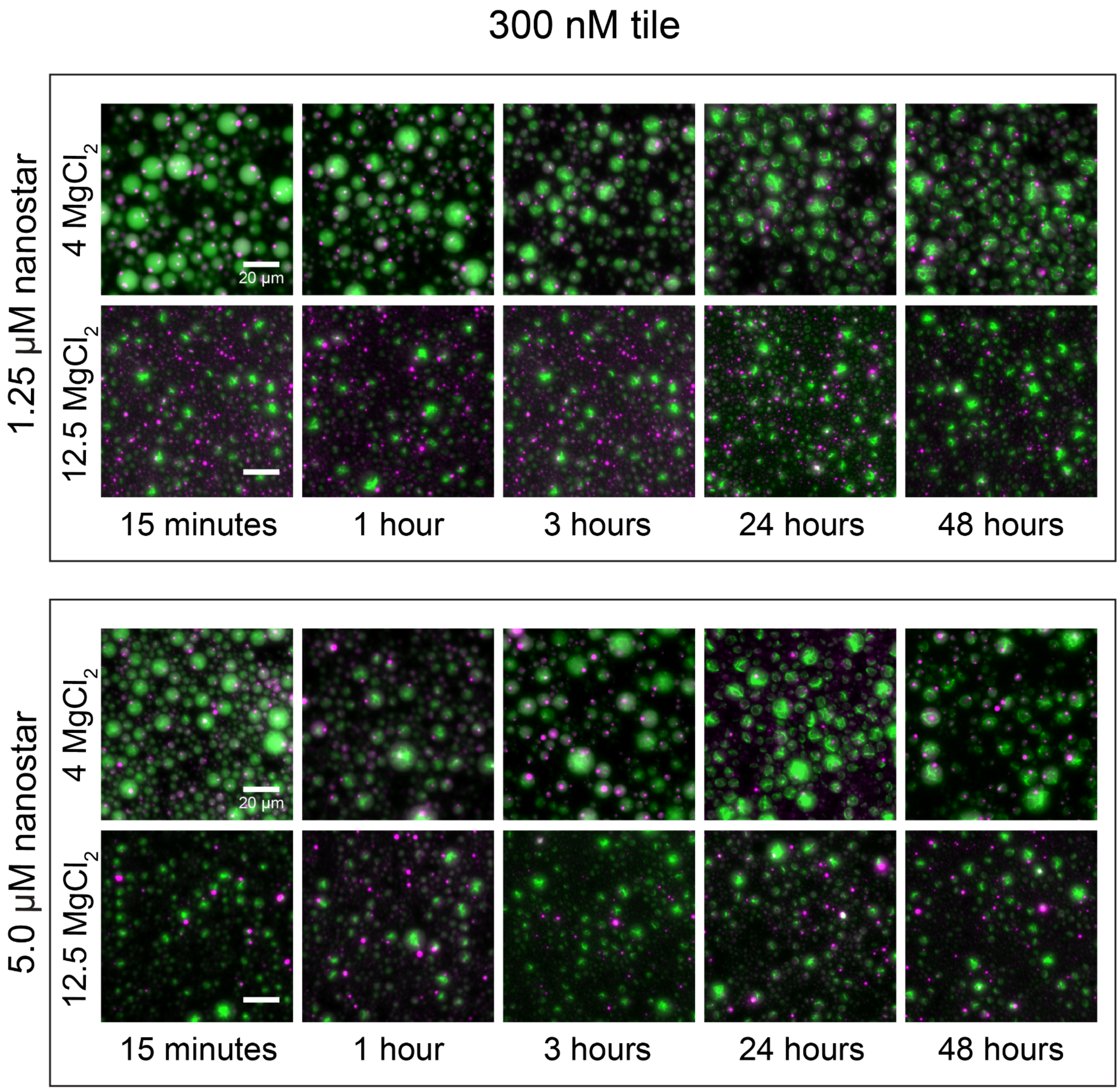


***Figure S4. Kinetic images of water-in-oil droplets show growth of condensates and nanotubes at 300 nM tile.*** *Increasing tile concentration to 300 nM facilitates earlier formation and saturation of nanotube networks at 4 mM $MgCl_2$ without forming nanotube aggregates. 12.5 mM $MgCl_2$ causes more significant aggregation of tiles at later timepoints.*

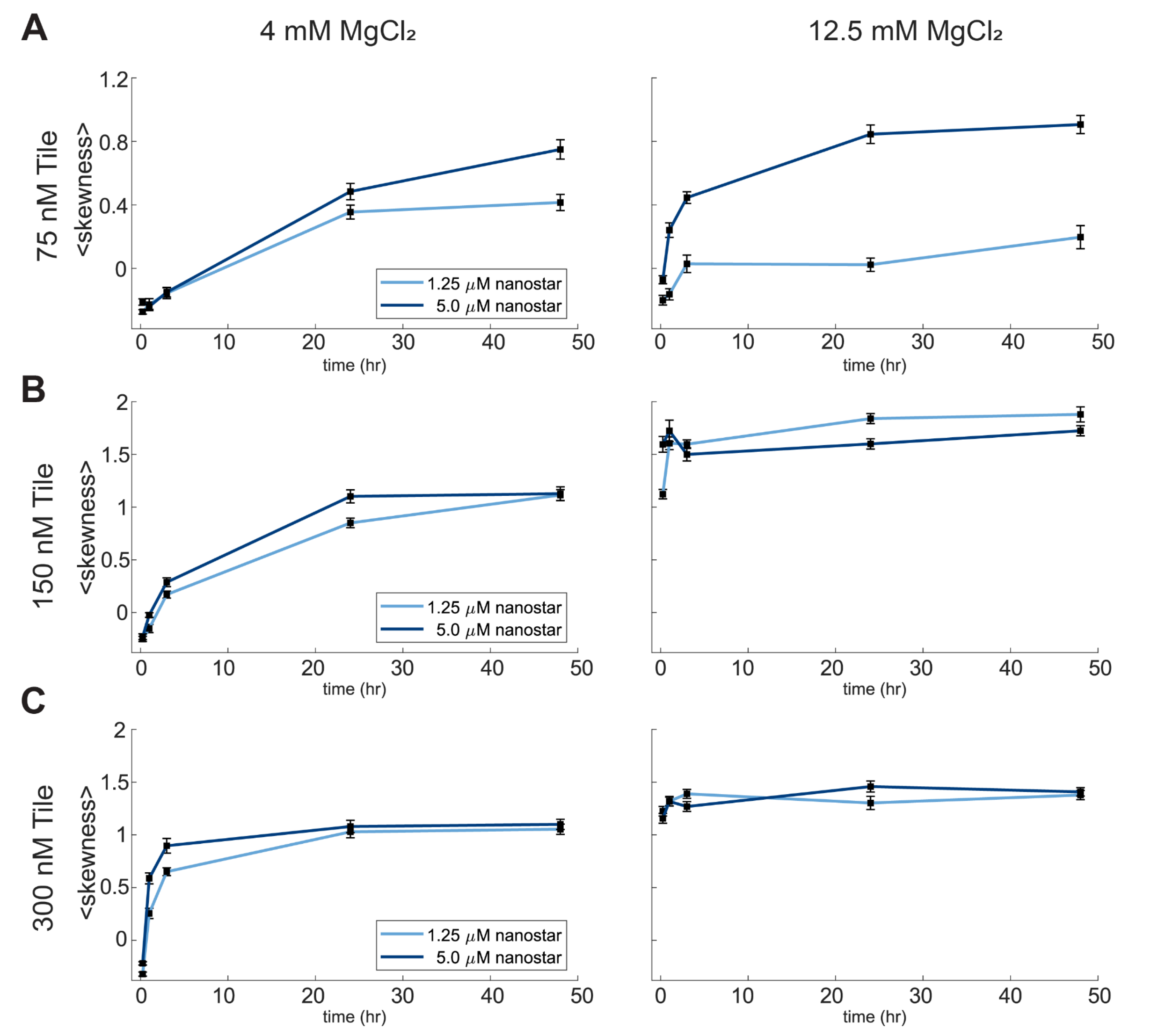


***Figure S5. Skewness plots for uncaged data show growth and aggregation of nanotubes up to 48 hours after water-in-oil droplet emulsion.*** *Skewness measures the distribution of FAM (tile) channel for circular ROIs, with increased skewness indicating more aggregation of nanotubes over time. Increasing concentration of tiles and $MgCl_2$ facilitates faster saturation of nanotube networks.*

.

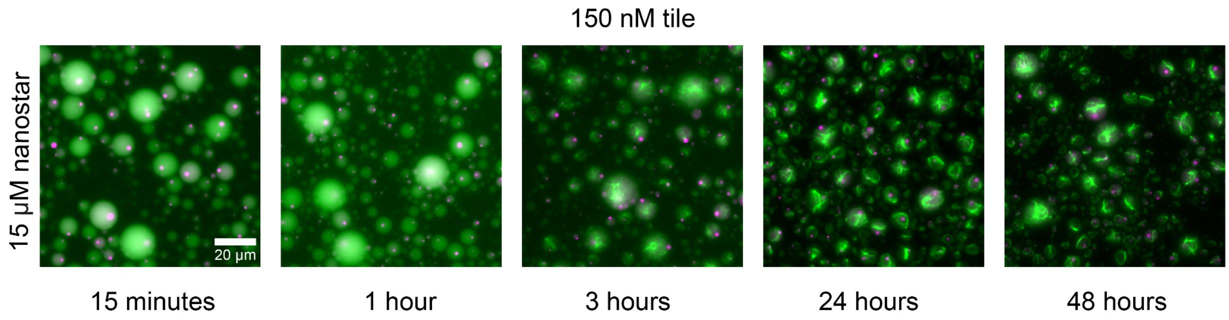


***Figure S6. Kinetic images of water-in-oil droplets at 15 µM nanostar and 150 nM tile concentration.***

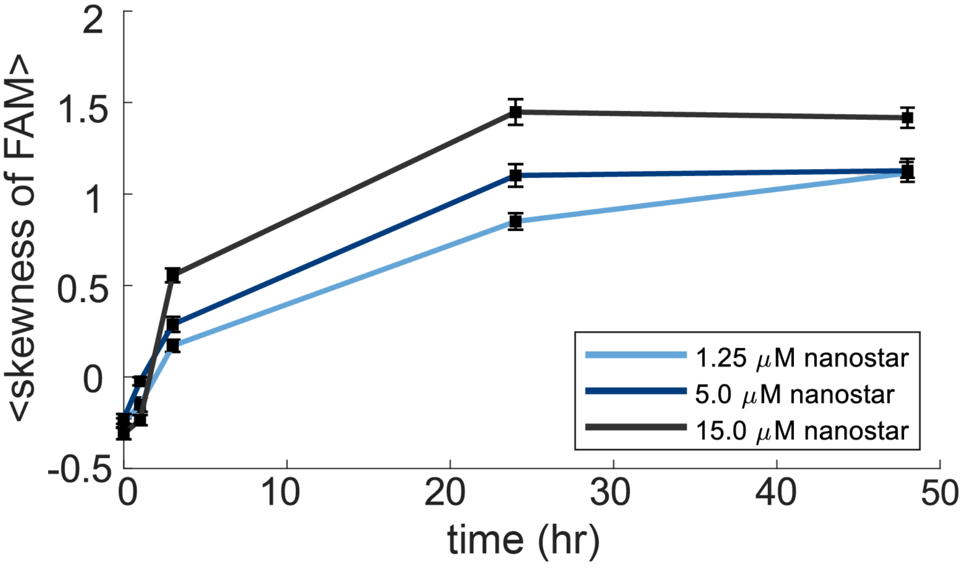


***Figure S7. Skewness plots comparing nanostar concentration at 150 nM tile shows increased aggregation of nanotubes with increased nanostar concentration.***

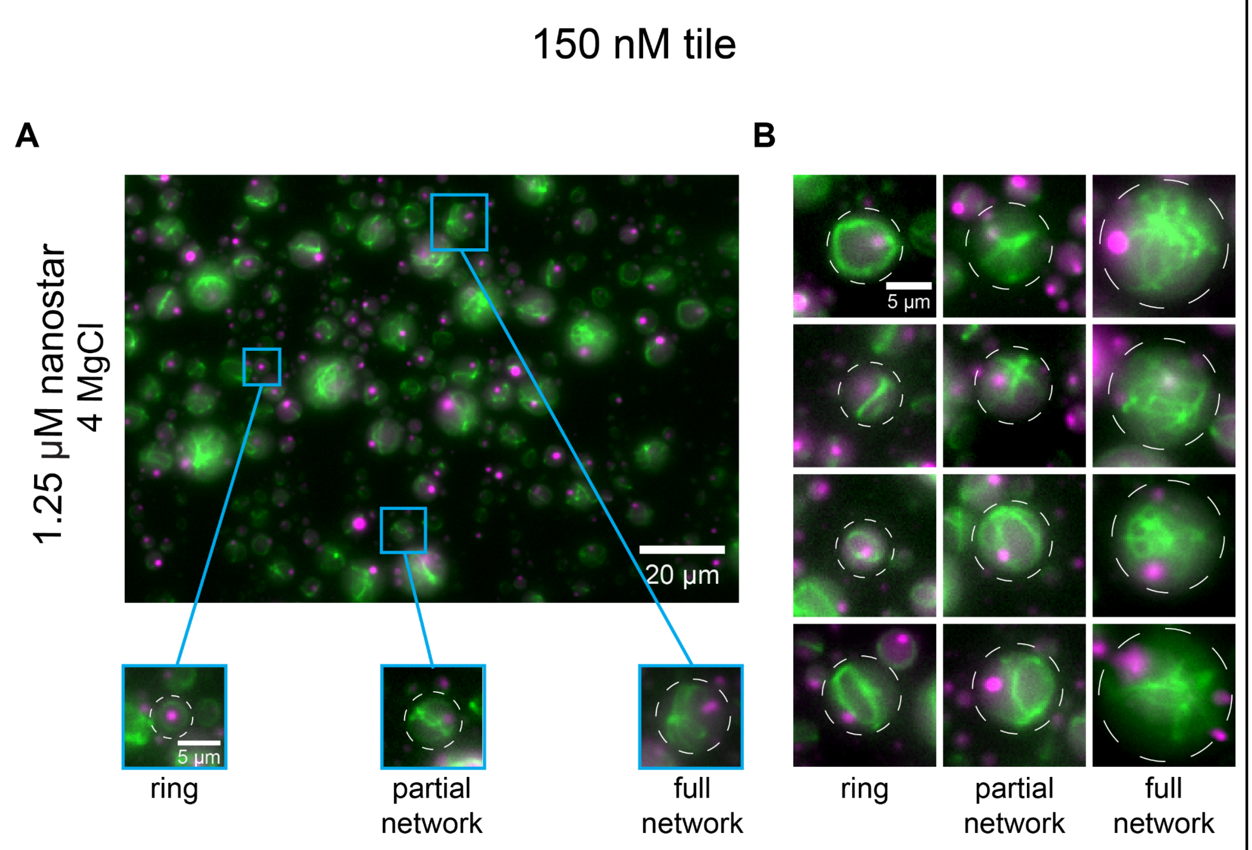


***Figure S8. Characterization of nanotube morphologies in water-in-oil droplets. A)*** *In samples of polydispersed water-in-oil droplets, nanotubes form distinct morphologies based on droplet diameter.* ***B)*** *Rings encircling the droplet form at the lowest droplet size and induce more crossover and branching of nanotubes as diameter increases, from partial networks (those with two or fewer nodes) to full networks (those with greater than two nodes).*

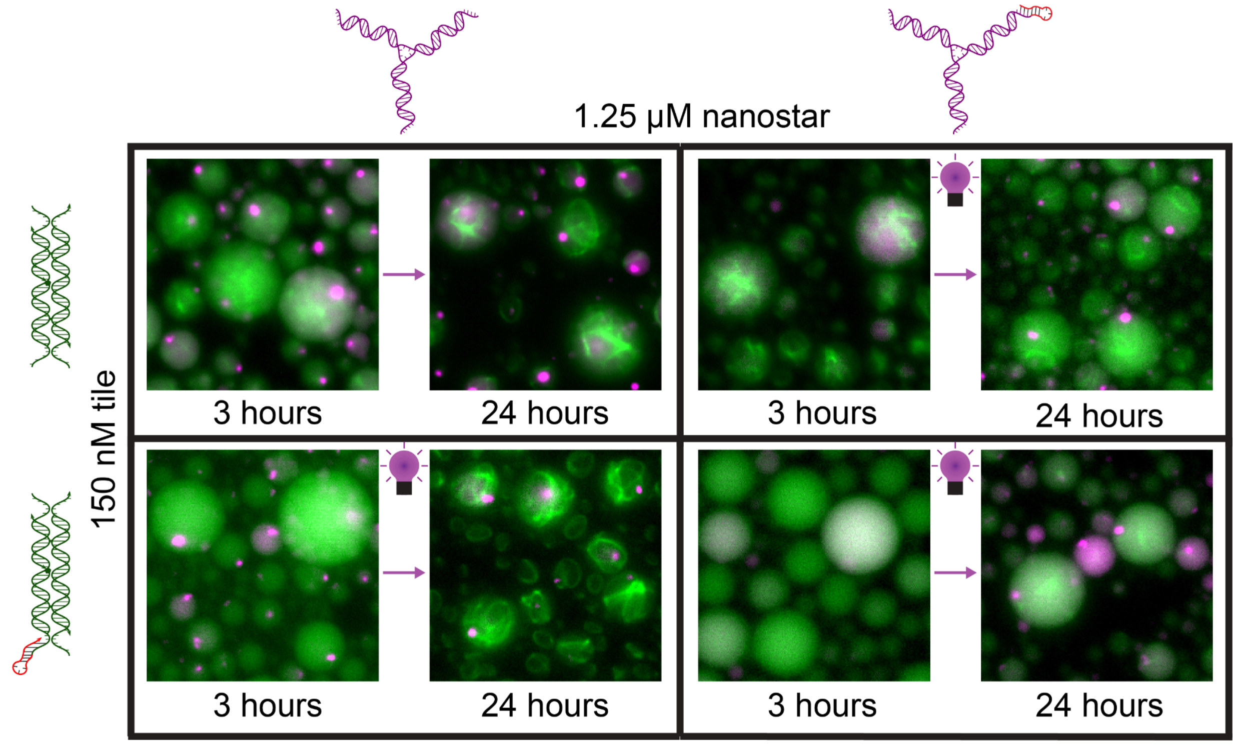


***Figure S9. Epifluorescence microscopy images of all permutations of caged nanostars and caged tiles show latent formation of either nanostructure.*** *From 3 hours after droplet emulsion to 24 hours, use of the photocleavable hairpin can prevent assembly of condensates and nanotubes until after irradiation of UV light. When both monomers are uncaged, formation of either structure occurs normally (top left), whereas caging both monomers allows for proper condensate assembly and significantly hampers nanotube assembly (bottom right).*

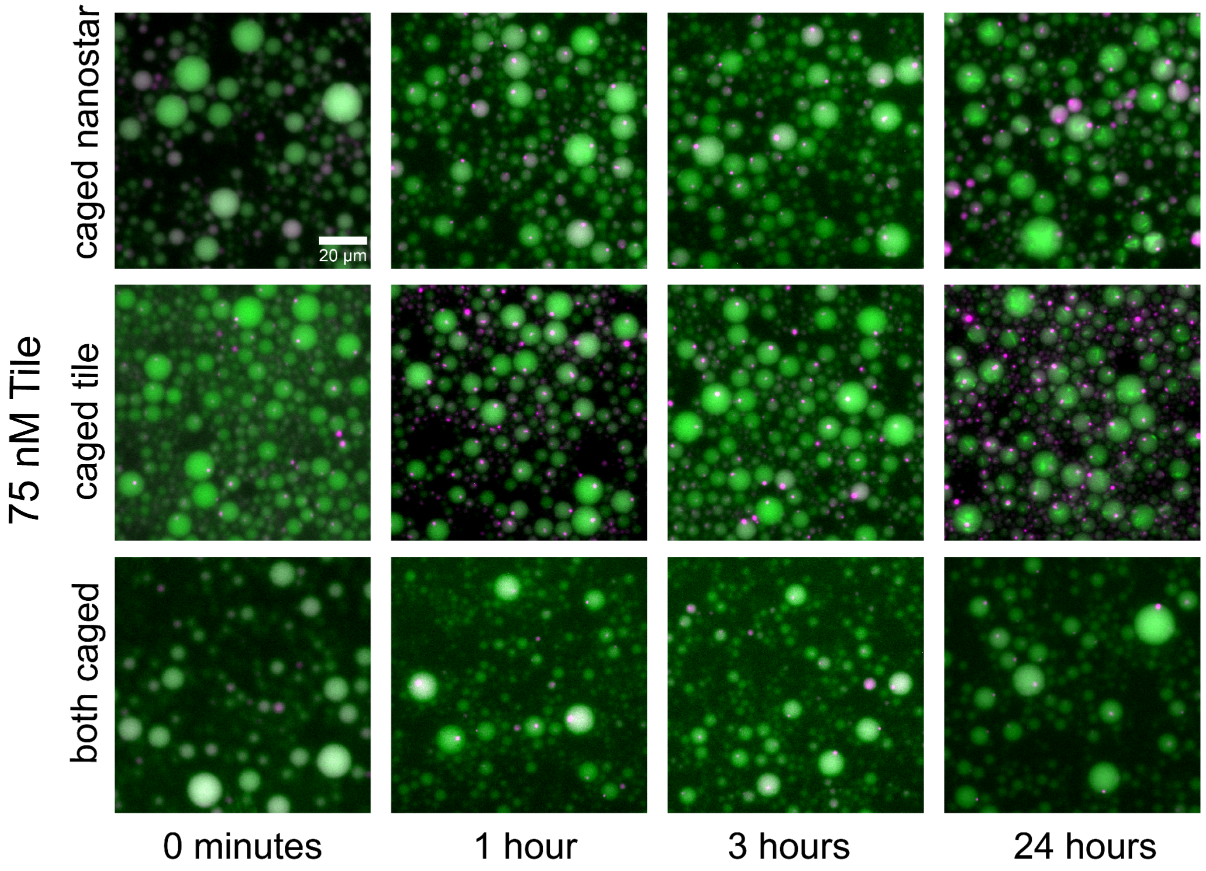


***Figure S10. Kinetic images show latent formation of condensates and nanotubes from monomers with a photocleavable hairpin after irradiation of UV light at 75 nM tile.*** *Occluding tile and nanostar assembly prevents formation of condensates and nanotubes for 3 hours until cleavage from UV irradiation (0 minute timepoint). Post-irradiation images show phase separation of condensates and growth of nanotubes up to 24 hours.*

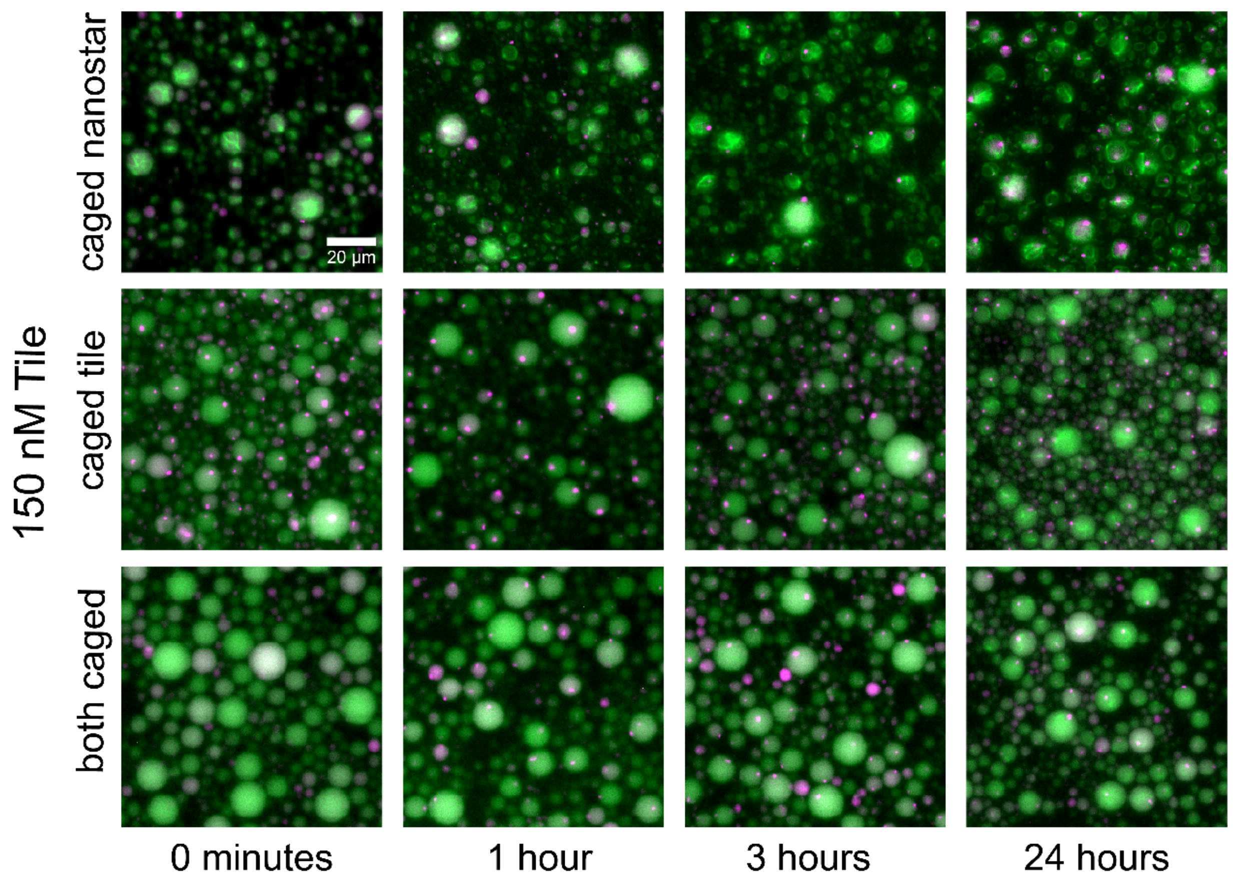


***Figure S11. Kinetic images show latent formation of condensates and nanotubes from monomers with a photocleavable hairpin after irradiation of UV light at 150 nM tile.*** *Condensate assembly occurs rapidly after UV irradiation on the sample. Nanotube growth is negatively affected by the presence of the dissociated hairpin domain in the caged tile condition in comparison to the caged nanostar condition. Caging both tiles and nanostars compounds this effect and disrupts aggregation of nanotubes up to 24 hours post-irradiation.*

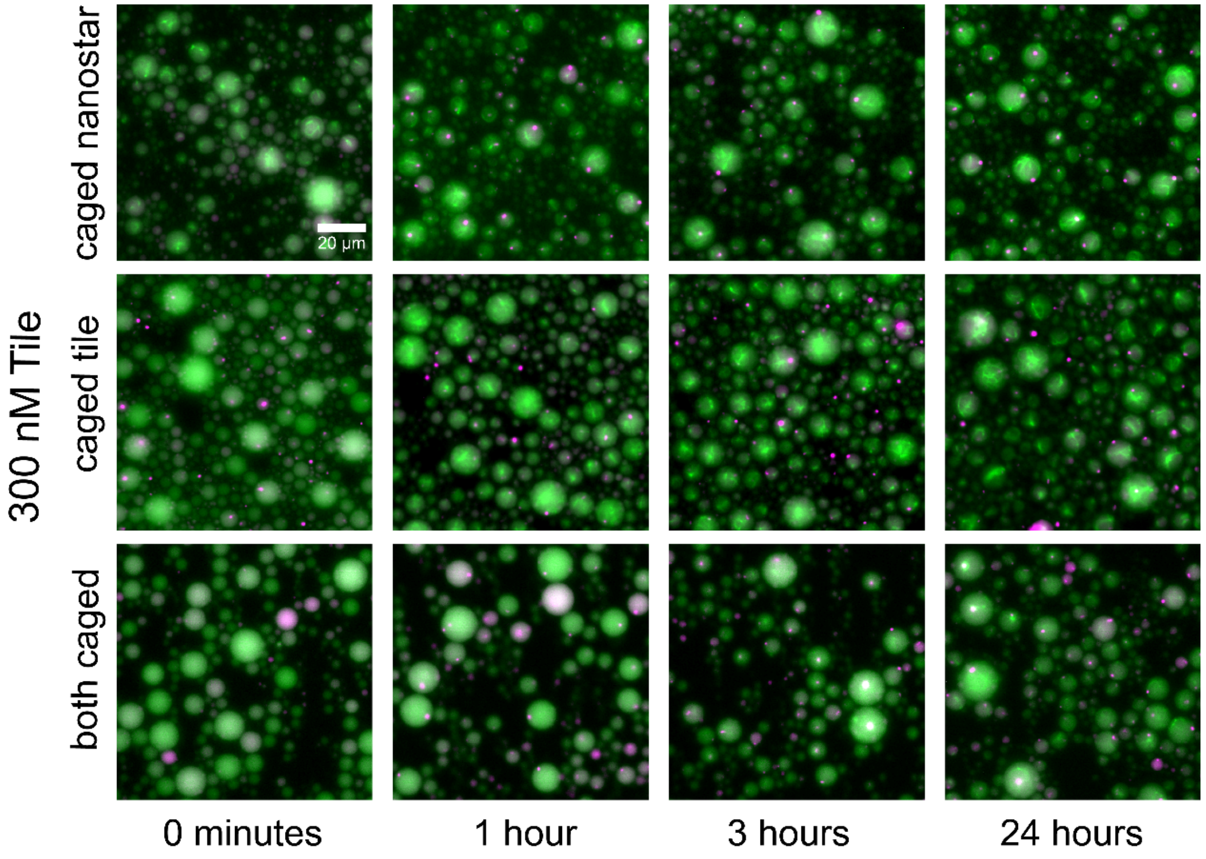


***Figure S12. Kinetic images show latent formation of condensates and nanotubes from monomers with a photocleavable hairpin after irradiation of UV light at 300 nM tile.*** *Condensate assembly occurs rapidly after UV irradiation on the sample. Nanotube growth is negatively affected by the presence of the dissociated hairpin domain in the caged tile condition in comparison to the caged nanostar condition. Caging both tiles and nanostars compounds this effect and disrupts aggregation of nanotubes up to 24 hours post-irradiation.*

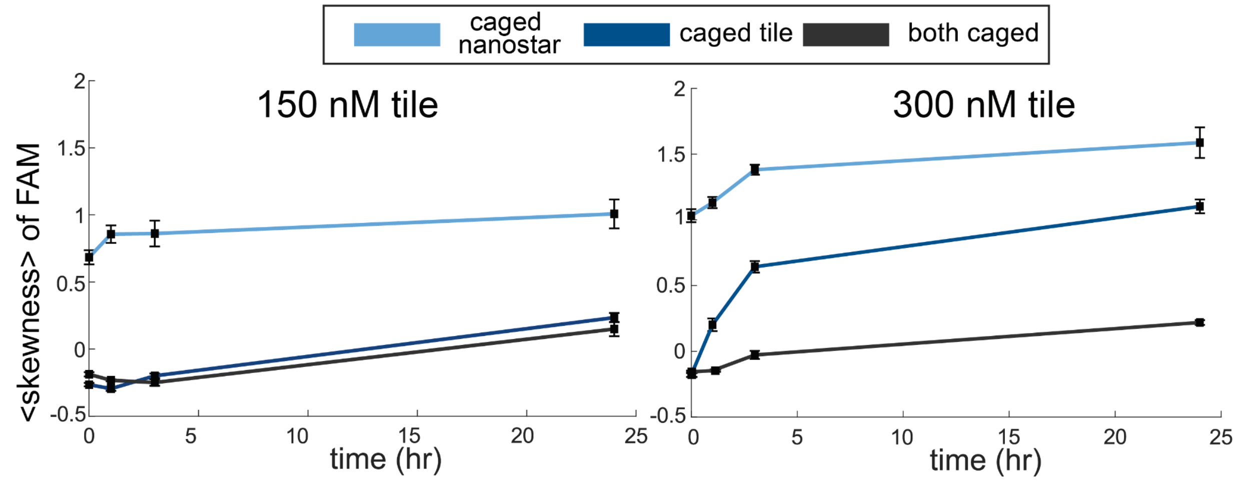


***Figure S13. Skewness plots demonstrate differences in the growth of nanotubes in water-in-oil droplets due to the confined presence of the photocleavable hairpin.*** *Skewness curves of the FAM (tile) channel show growth of nanotubes over time, though caging the tiles and caging both tiles and nanostars affects the aggregation of nanotubes in confinement. At 0 minutes, conditions employing the caged tile show no nanotube formation, while the caged nanostar has higher skewness values, demonstrating nanotube growth for 3 hours prior to UV irradiation with uncaged tiles.*

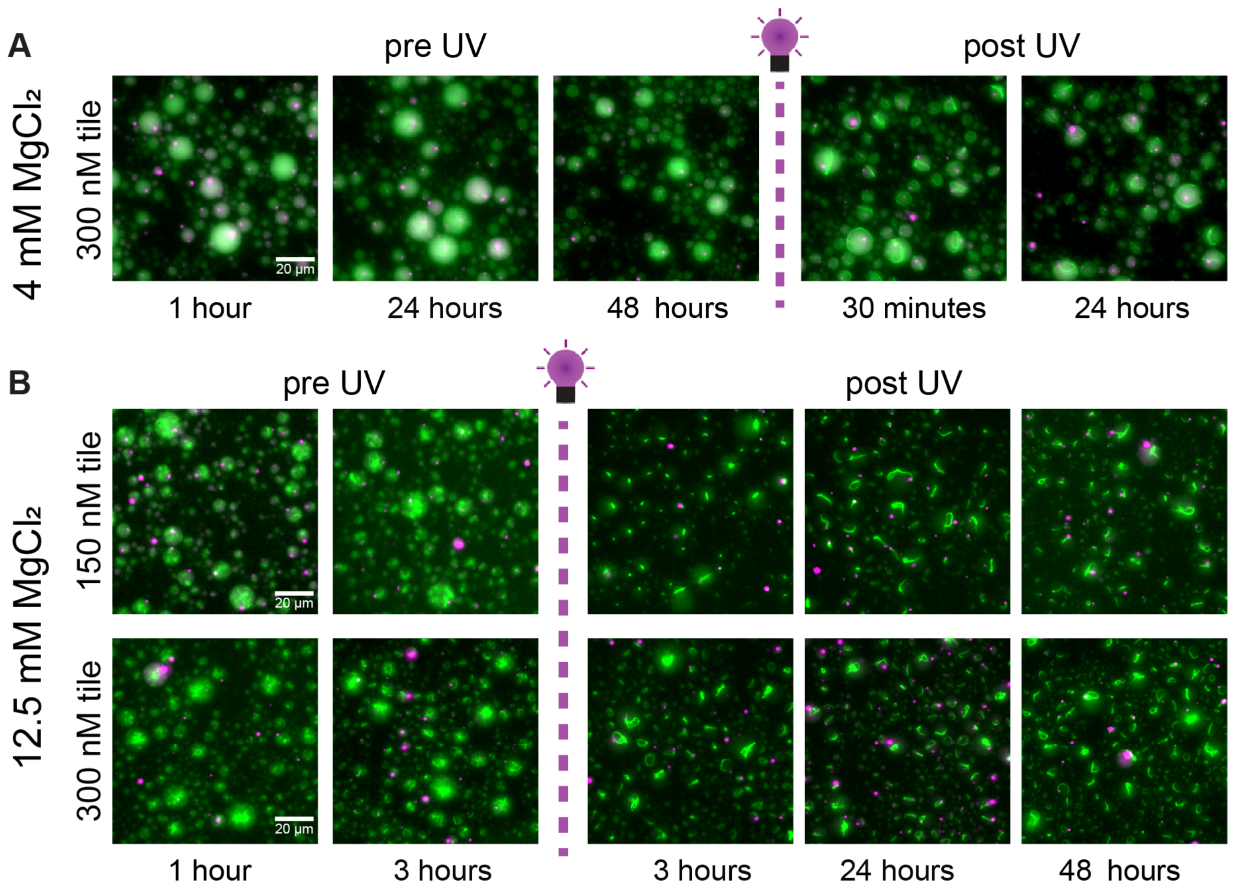


***Figure S14. Usage of a photocleavable linker domain allows full nanotube networks to form, then induce restructuring of the nanotube network into ring-like structures upon irradiation of UV light.*** *A palindromic linker domain is added to the end of one tile sticky end, which facilitates nanotube bundling into rings in water-in-oil droplets.* ***A)*** *At 4 mM $MgCl_2$, the bulky photocleavable linker hairpin partially blocks sticky end interactions between tiles. After 48 hours of nanotube growth, UV irradiation induces bundling of nanotubes and ring formation within 30 minutes, and ring morphology is retained up to 24 hours post-irradiation.* ***B)*** *At 12.5 mM $MgCl_2$, nanotube growth is rapid and forms full networks in 3 hours. UV irradiation facilitates nanotube bundling into rings, rather than full networks and aggregates at 3 hours and retains ring morphology up to 48 hours post-irradiation.*

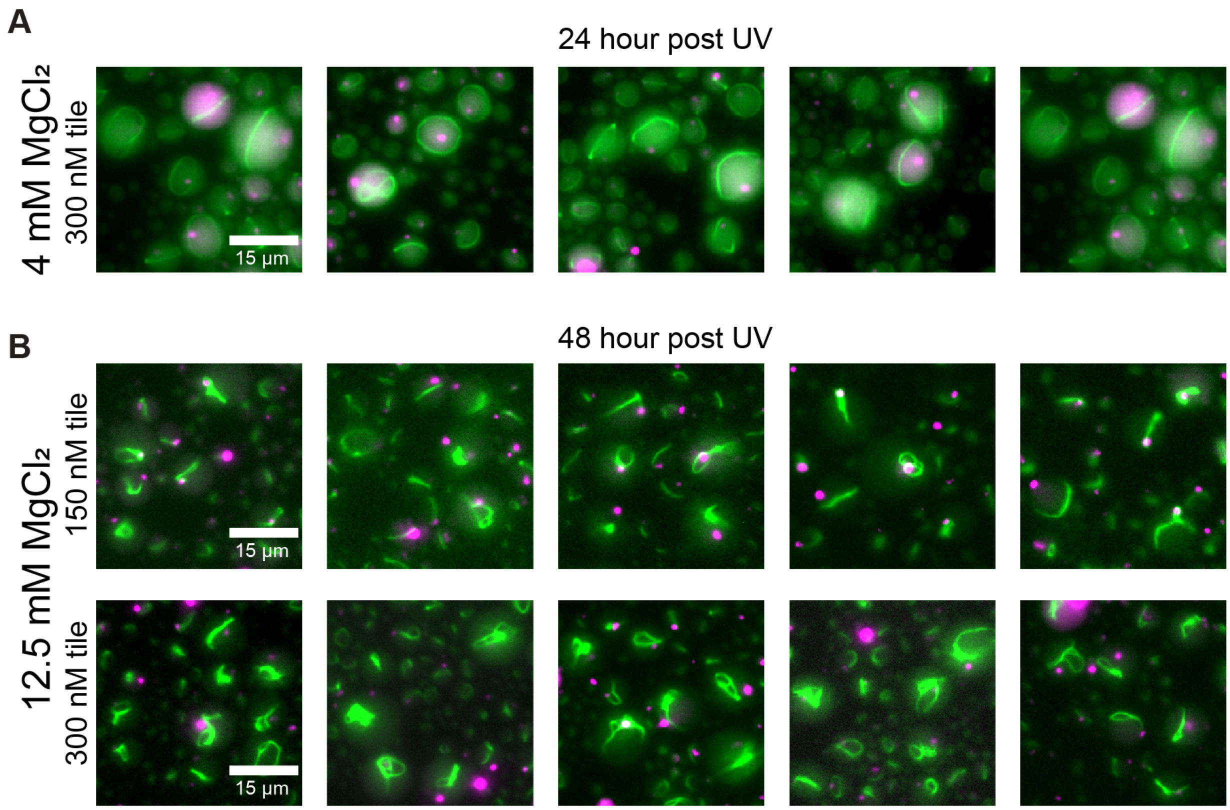


***Figure S15. Epifluorescence microscopy images depicting nanotube bundling into rings at 24 or 48 hours post-irradiation of UV light. A)*** *At 4 mM $MgCl_2$, tiles bundle into rings that encircle the full diameter of the water-in-oil droplet due to increased mobility of individual tiles prior to full nanotube network assembly.* ***B)*** *At 12.5 mM $MgCl_2$, nanotubes retain full network morphology and begin bundling into rings, but with significant branching off fully formed rings.*

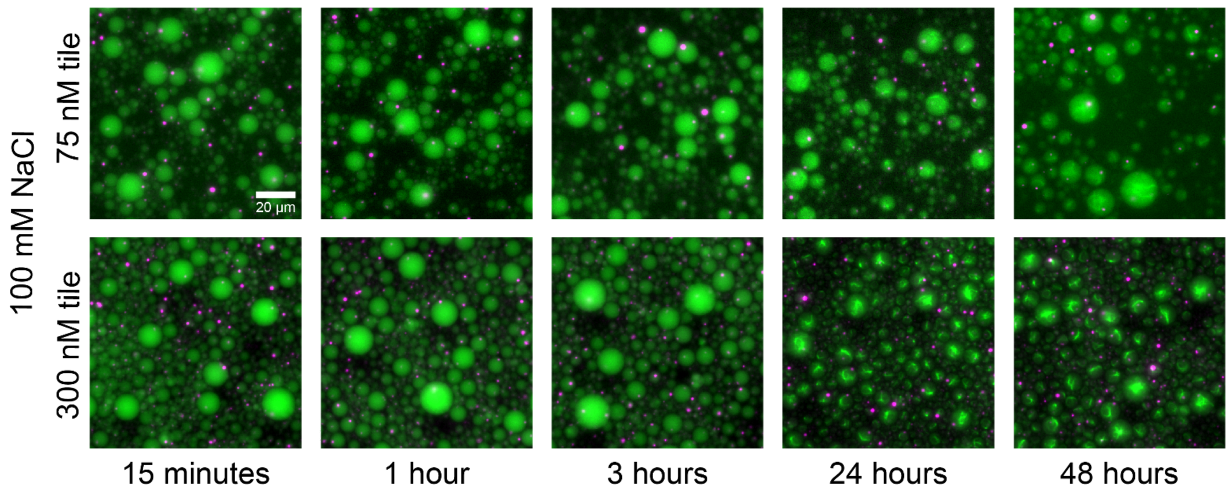


***Figure S16. Kinetic images of isothermal assembly and growth of condensates and nanotubes in water-in-oil droplets with TANa buffer.*** *Assembly of condensates inside water-in-oil droplets with TANa buffer is significantly slower at 75 nM tile because low concentration provides a high energetic barrier to initial assembly of tiles compared to 300 nM. Nanotube networks form properly at 300 nM tile at 24 hours and retain full network morphology at up to 48 hours after droplet emulsion.*

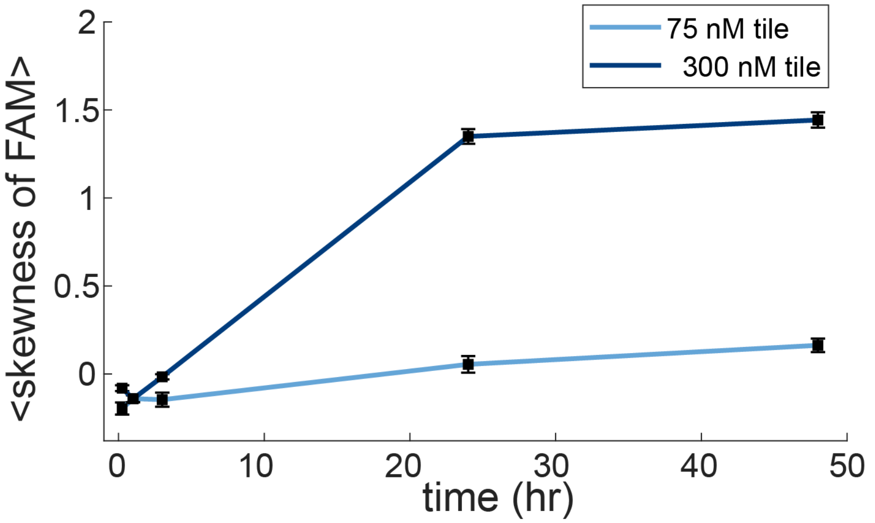


***Figure S17. Skewness plots of different concentrations of tile in water-in-oil droplets using TANa buffer.*** *Due to the isothermal formation of tiles prior to nanotube assembly, nanotubes show little growth at 75 nM and grow regularly in the 300 nM condition.*

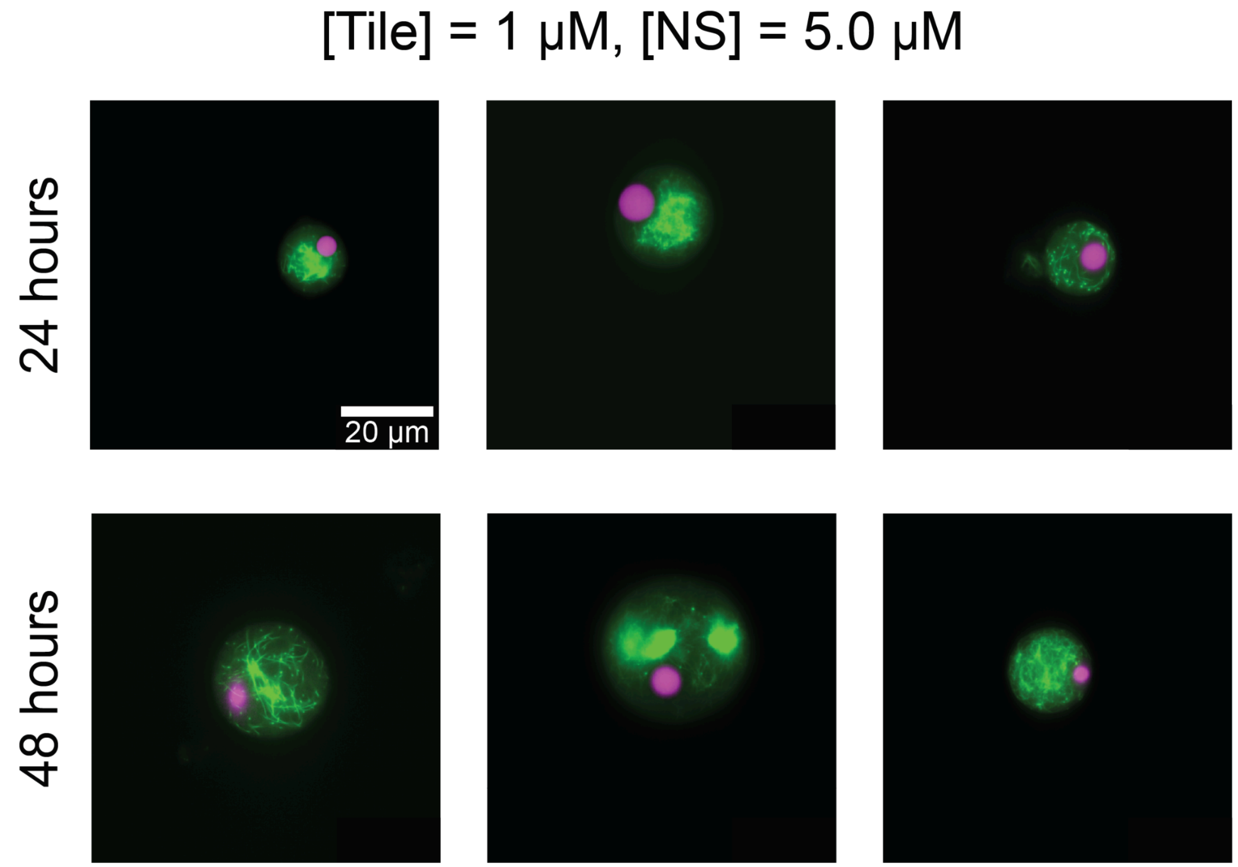


***Figure S18. Epifluorescence microscopy images of nanotubes and condensates confined inside GUVs after isothermal formation with 100 mM NaCl TANa buffer.*** *TANa buffer uses monovalent NaCl to facilitate formation of tiles and nanostars prior to assembly into condensates and nanotubes. Increasing the tile concentration to 1 µM causes rapid formation of nanotube networks, but often causes aggregation of nanotubes instead of full networks inside GUVs.*
